# Prevention of chromatin destabilization by FACT is crucial for malignant transformation

**DOI:** 10.1101/499376

**Authors:** Poorva Sandlesh, Alfiya Safina, Imon Goswami, Laura Prendergust, Spenser Rosario, Eduardo C Gomez, Jianmin Wang, Katerina V Gurova

**Affiliations:** Department of Cell Stress Biology, Roswell Park Comprehensive Cancer Center, Carlton and Elm Streets, Buffalo, NY, 14127; Department of Cancer Genetics, Roswell Park Comprehensive Cancer Center, Carlton and Elm Streets, Buffalo, NY, 14127; Department of Bioinformatics, Roswell Park Comprehensive Cancer Center, Carlton and Elm Streets, Buffalo, NY, 14127

**Author notes:** present address: – Division of Reproductive Science in Medicine, Department of Obstetrics and Gynecology, Robert H. Lurie Comprehensive Cancer Center, Feinberg School of Medicine, Northwestern University, Chicago, IL 60611. Corresponding author: Katerina Gurova, Department of Cell Stress Biology, Roswell Park Comprehensive Cancer Center, Carlton and Elm Streets, Buffalo, NY, 14127.

## Abstract

Expression of histone chaperone FACT is increased in tumors and associated with poor prognosis. We investigated why aggressive tumor cells need FACT using a model where FACT could be turned off and confirmed that while FACT is not essential for non-tumor cells, cells become dependent on FACT following oncogene-induced transformation. We compared the phenotypic and transcriptional changes induced by FACT loss and excluded a direct role for FACT in the transcription of genes essential for the viability of transformed cells. Moreover, we established that in immortalized and transformed cells, FACT has a weak negative effect on gene expression. At the same time, we observed a positive correlation between FACT enrichment and the rate of transcription, which was consistent with previous reports. To explain these puzzling observations, we hypothesized that FACT does not facilitate transcription elongation in transformed cells, but prevents nucleosome loss associated with transcription. Indeed, we observed destabilization of chromatin in immortalized and transformed cells upon FACT loss. Furthermore, transformed cells had less stable chromatin than non-transformed cells, which made them vulnerable to FACT loss. However, the mechanisms of cell death upon chromatin destabilization needs to be established. Our data suggest that malignant transformation is accompanied by general chromatin destabilization, and FACT prevents irredeemable chromatin loss.

## Introduction

Multiple chromatin alterations are found in cancer, such as mutations and changes in the expression of histones, chromatin remodeling factors, histone chaperones, and enzymes that post-translationally modify histones (reviewed in (Morgan and Shilatifard 2015; Ferraro 2016; Flavahan et al. 2017)). The benefits that these alterations provide to tumor cells are unclear. The prevailing hypothesis is that these alterations lead to changes in the expression of genes that promote cell growth or inhibit differentiation and cell death. However, this hypothesis is not completely satisfying because it does not explain how chromatin alterations (e.g., mutations in core histones) with extensive genome-wide effects can lead to changes in the transcription of specific genes that are beneficial for cancer cells.

One example of such an alteration is the frequent overexpression of histone chaperone FACT (Facilitates Chromatin Transcription) in multiple human tumors (Garcia et al. 2013; Carter et al. 2015; Ding et al. 2016; Fleyshman et al. 2017). FACT consists of two subunits in higher eukaryotes: Structure Specific Recognition Protein 1 (SSRP1) and Suppressor of Ty 16 (SPT16). Both subunits are highly conserved in all eukaryotes and perform similar functions. They interact with all components of the nucleosome (i.e., histone oligomers and DNA) and are involved in replication, transcription, and DNA repair (reviewed in (Gurova et al. 2018)). FACT is not only overexpressed in different types of tumors but is also associated with poor prognosis (Garcia et al. 2013; Carter et al. 2015; Dermawan et al. 2016; Attwood et al. 2017). Moreover, genetic or chemical inhibition of FACT has strong anti-cancer effects in multiple cancer models (Gasparian et al. 2011; Garcia et al. 2013; Carter et al. 2015; Dermawan et al. 2016). At the same time, mammalian FACT is not expressed or is expressed at much lower levels in non-tumor cells *in vitro* and differentiated cells *in vivo* (Garcia et al. 2011). Inhibition of FACT in FACT-positive normal cells has little effect on cell growth or viability (Garcia et al. 2013; Mylonas and Tessarz 2018), (Kolundzic et al. 2018). These findings suggest that FACT may be a promising target for anti-cancer treatment. However, how FACT supports the viability of tumor cells is unclear.

In cell-free experiments, FACT was essential for transcription elongation through nucleosomal DNA (Orphanides et al. 1998; Orphanides et al. 1999). Based on these data, when we first noticed that FACT was enriched at coding regions of so-called “pro-cancerous genes” (i.e., genes involved in cell proliferation, response to stress, and maintenance of pluripotency) (Garcia et al. 2013), we assumed that FACT was involved in the transcription elongation of these genes, many of which are essential for tumor growth. However, there were many unclear issues with this interpretation, including how FACT selected these genes because FACT does not have sequence-specific DNA binding or histone modification “reader” domains. If the elongating RNA polymerases recruited FACT, then why would its inhibition be much more toxic for tumor than normal cells? Furthermore, depletion of FACT from tumor cells, which were the most sensitive to FACT knockdown, did not result in the inhibition of the expression of these genes (Fleyshman et al. 2017). Similarly, it was recently shown that there was no inhibition of the transcription of FACT-enriched genes in mouse embryonic stem cells or human fibroblasts (Kolundzic et al. 2018; Mylonas and Tessarz 2018).

Several groups recently reported that mammalian FACT could not bind the folded nucleosome (Tsunaka et al. 2016; Safina et al. 2017; Wang et al. 2018), which makes it difficult to explain how FACT can remove the nucleosomal barrier for transcription and replication. However, FACT can bind destabilized nucleosomes or nucleosomes with unwrapped DNA, which was shown in cells and cell-free systems (Tsunaka et al. 2016; Safina et al. 2017; Wang et al. 2018). In addition, several previous reports showed that mutation of the FACT subunits in yeast or depletion of the subunits from human HeLa cells was accompanied by histone loss from transcribed regions (Morillo-Huesca et al. 2010; Myers et al. 2011; Carvalho et al. 2013; Erkina and Erkine 2015; Feng et al. 2016). These findings suggest that major function of FACT may not be disassembly of nucleosomes during transcription, but in opposite, prevention of loss of histones, when their contacts with DNA are weakened, e.g during RNA or DNA polymerases passage (Gurova et al. 2018)). This model explains most cell-based observations but not why FACT is essential for aggressive tumor cells.

In this study, we used an isogenic cell model of step-by-step malignant transformation with a conditional knockout of the *Ssrp1* gene. Using this model, we evaluated the phenotypic changes associated with transcription, replication, and chromatin organization upon FACT loss in matched primary, immortalized, and transformed cells.

## Results

### 1. Development of conditional *Ssrp1* knockout cell model with different basal levels of FACT

Previously, we observed that tumor cells express higher levels of the FACT subunits, and their viability is more dependent on FACT expression than primary or immortalized non-tumor cells (Garcia et al. 2011; Garcia et al. 2013), suggesting that FACT function is vital for tumor cells. To understand this difference in FACT dependency, we generated isogenic cells from mouse skin fibroblasts (MSFs) isolated from *Ssrp1^fl/fl^* CreER^T2+/+^ mice, in which the *Ssrp1* gene can be deleted by tamoxifen treatment (Sandlesh et al. 2018). As a negative control, we used cells from *Ssrp1^fl/+^* CreER^T2+/+^ mice because deletion of one allele of *Ssrp1* did not affect the mouse phenotype (Cao et al. 2003). We previously demonstrated that depletion of SSRP1 leads to an efficient and rapid loss of both SSRP1 and SPT16 proteins (Safina et al. 2013). Thus, the whole FACT complex can be eliminated from these cells by the administration of the active metabolite of tamoxifen, 4-hydroxytamoxifen (4-OHT).

Primary MSFs are highly sensitive to contact inhibition, survive in culture for 4 to 5 passages, and then undergo replicative senescence. The MSFs were transduced with the genetic suppressor element (GSE) 56, an inhibitor of tumor suppressor p53 (Ossovskaya et al. 1996). MSF-GSE56 cells became immortal but were still sensitive to contact inhibition (Fig. 1A), did not grow in semisolid medium and did not form tumors in mice. MSF-GSE56 cells were subsequently transduced with the mutant H-Ras^V12^ oncogene. These cells (MSF-GSE56-HRas) lost contact inhibition and formed foci in dense culture (Fig. 1A). They also grew in semisolid medium and quickly developed aggressive tumors in mice (Fig. 1B-D), i.e., acquired a fully transformed phenotype. The primary MSFs had low but detectable levels of SSRP1 and SPT16, which were elevated in immortalized MSF-GSE56 cells and further increased in the transformed MSF-GSE56-HRas cells (Fig. 1A). In all experiments, we used primary cells isolated from 2 to 4 individual mice as biological replicates, which were immortalized and then transformed as described above. The main figures include the mean data for all tested cell variants or representative images. Data for additional cell variants are available in the supplementary materials. Primary, immortalized and transformed cells from the same mouse are labeled with the same number.

**Figure 1.**
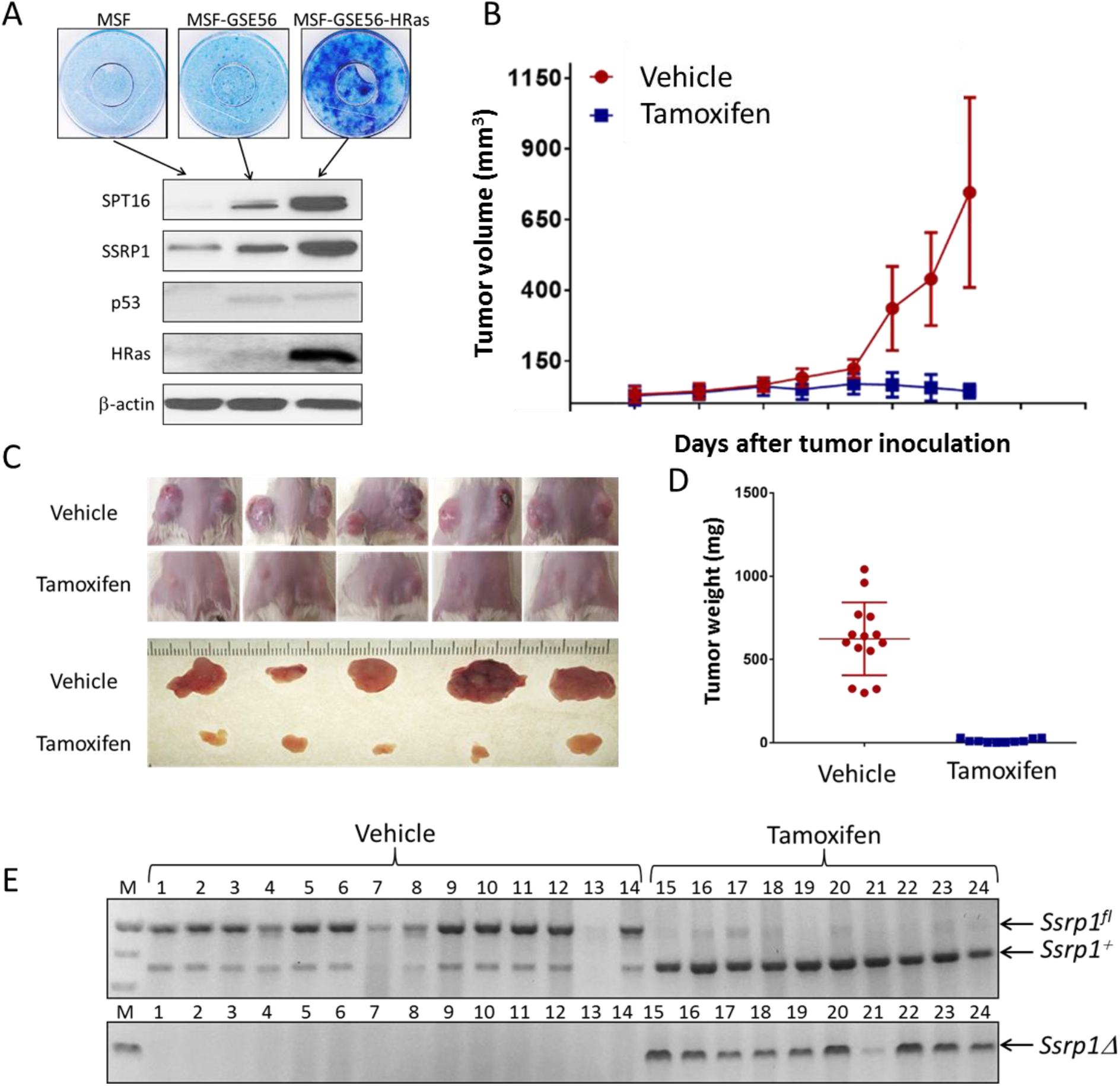
Model of *in vitro* transformation. A. Primary (Pr) mouse skin fibroblasts (MSF) were isolated from the tail tips of *Ssrp1^fl/fl^;CreERT^2^+/+* mice and transduced with GSE56 to become immortalized (Im) or GSE56 and HRas^V12^ oncogene to become transformed (Tr). Top – methylene blue stained plates of Pr, Im, and Tr cells at confluency. Bottom – western blotting of lysates from the three cell types probed with the indicated antibodies. B. Growth of Tr cells in SCID mice treated with tamoxifen or vehicle (control n = 13, tamoxifen n = 14). Data are presented as the mean tumor volume ± SD. C. Photographs of mice and tumors from B. D. Weight of tumors from B at the end of experiment. E. Testing of excision of *Ssrp1* (appearance of *Ssrp1D*) in tumors at the end of treatment with tamoxifen using PCR of genomic DNA extracted from individual tumors. *Ssrp1^fl^*, mutant allele; *Ssrp1^+^*, wild-type *Ssrp1; Ssrp1Δ*, excised allele. *Ssrp1^+^* is present in tamoxifen-treated tumors most likely due to its presence in stroma.

### 2. Different consequences of FACT depletion in primary, immortalized and transformed cells

4-OHT administration resulted in the disappearance of *Ssrp1* mRNA and corresponding protein between days 3 and 5 after the start of treatment (Fig. 2A and S1). SPT16 protein levels decreased with the same kinetics. *Supt16* (i.e., the gene encoding SPT16 in mice) mRNA remained unchanged (Fig. S1). Treatment of primary cells with 4-OHT did not affect their growth. In contrast, the same treatment significantly inhibited the growth of the transformed cells and, to a lesser extent, the immortalized cells (Fig. 2B and S2). Importantly, tumors that formed from the transformed cells in SCID mice quickly disappeared after the administration of tamoxifen (Fig. 1B-E).

**Figure 2.**
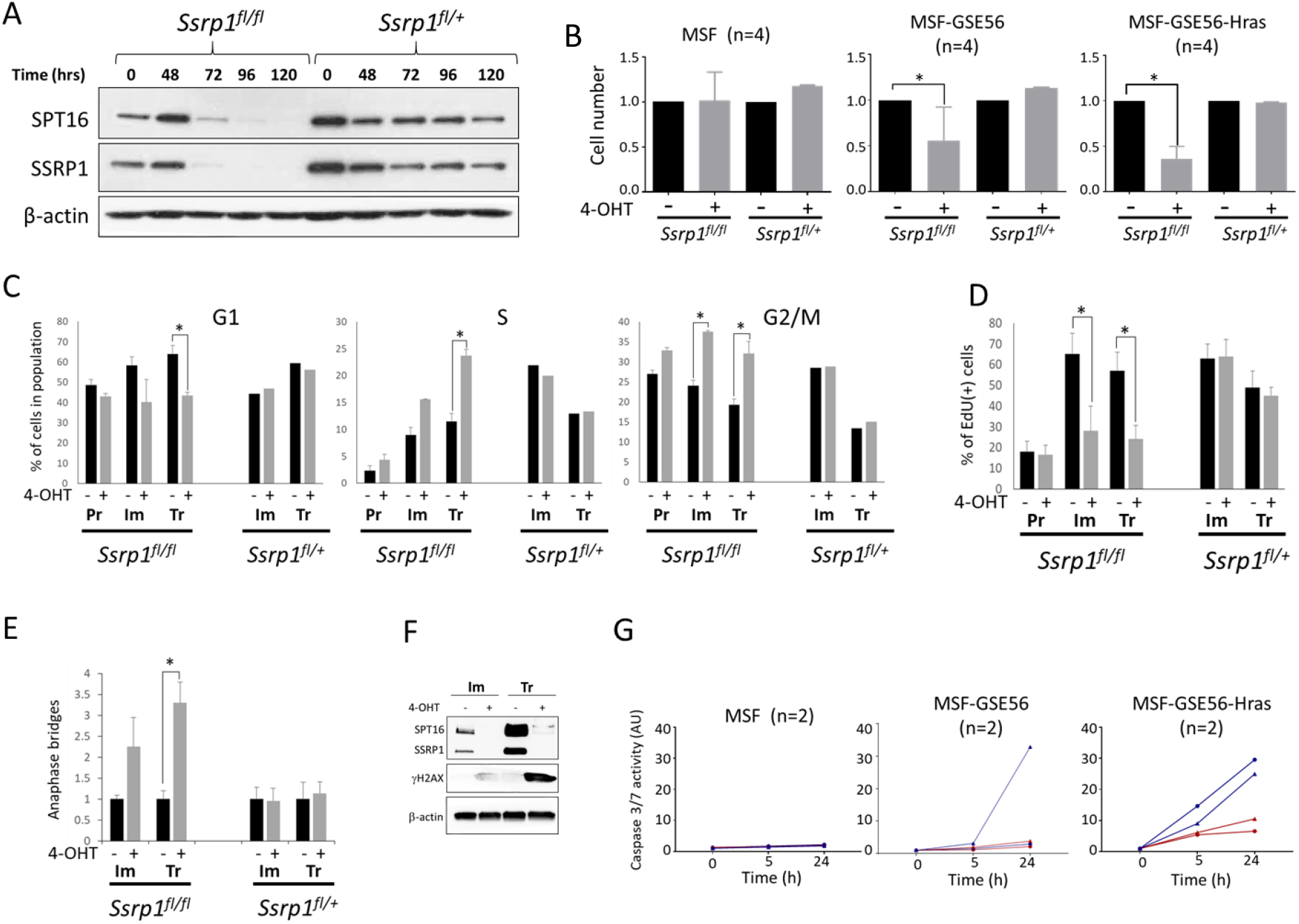
Effect of 4-OHT administration on primary (Pr), immortalized (Im), and transformed (Tr) cells established from *Ssrp1^fl/fl^;CreERT^2+/+^ or Ssrp1^fl/+^;CreERT^2+/+^* mice. A. Excision of *Ssrp1^fl/fl^* results in disappearance of SSRP1 and SPT16 proteins after 5 days of treatment with 4-OHT. Western blotting of extracts from Im cells at indicated time points after the start of treatment with 4-OHT. B. Viability of Pr, Im, and Tr cells with and without FACT. Equal numbers of each cell type were plated at the end of treatment with 4-OHT. Two days later, the relative number of cells was assessed using a resazurin-based cell viability assay. The bars represent the mean ± SD (n = 4) (see Fig. S2 for the data for the individual cell lines). C. Cell cycle distribution for cells replated following 4-OHT treatmen. Analysis was performed 24 h after replating. The bars represent the means ± SD (n = 2) (see Fig. S3 for the data from each individual cell line). D. EdU incorporation in cells treated with 4-OHT or vehicle. The bars represent th means ± SD (n = 2) (See Fig. S4 for the data from each individual cell line). E. Average number of anaphase bridges per number of mitoses. The bars represent th means ± SD (n = 2). Representative cell images are shown on Fig. S5. F. Accumulation of phosphorylated histone H2AX (gH2AX) and loss of FACT subunits in cells treated with 4-OHT or vehicle for five days. Western blotting of total cell lysates probed with the indicated antibodies. G. The activity of caspases 3 and 7 in the cells treated with 4-OHT (blue) or vehicle (red) measured at different time points after caspase substrate addition (n = 2 per genotype).

To understand the mechanism of the differential FACT-dependence of the three types of MSFs, we determined the changes in the cell cycle upon *Ssrp1* knockout (KO). The cell cycle distribution of the primary cells remained unchanged following 4-OHT treatment. In contrast, this treatment caused an accumulation of transformed and, to a lesser extent, immortalized cells in the S and G2/M phases of the cell cycle (Fig. 2C and S3). At the same time, DNA replication was reduced upon FACT depletion in these cells (Fig. 2D and S4). These data together with the cell viability data suggested that transformed and, to a lesser degree, immortalized cells have a problem moving through the S phase in the absence of FACT. The increased proportion of the cells in the G2/M phase may be due to the start of mitosis in cells with decreased S-phase checkpoint control as a result of p53 inactivation (i.e., when cells with unfinished replication are unable to separate DNA during mitosis). To test this hypothesis, we assessed the number of anaphase bridges present in the cells, which identifies cells undergoing mitosis with not-fully-replicated DNA, and found a significantly elevated number of anaphase bridges after FACT depletion in the transformed cells (Fig. 2F and S5). In immortalized cells, the same trend was observed, but it did not reach statistical significance. Consistent with this data, 4-OHT treatment resulted in the appearance of or increased levels of phosphorylated histone H2AX (γH2AX) in immortalized and transformed cells, respectively, which is indicative of DNA damage (Fig. 2E). We also detected activated caspase 3/7 after 4-OHT administration in transformed, and in one of three replicates of immortalized cells, but not in primary cells (Fig. 2G). 4-OHT did not cause any of these effects in cells obtained from mice heterozygous for *Ssrp1^fl^* allele (Fig. 2 and S1-S5).

The incorporation of 5-ethynyl-2′-deoxyuridine (EdU) into DNA was significantly lower in primary MSFs compared to the immortalized and transformed cells (Fig.2D and S4). The primary cells also had a much lower proportion of cells in S phase (Fig. S3). FACT has been shown to be essential for replication in frog, chicken, and human tumor cells (Okuhara et al. 1999; Tan et al. 2006; Abe et al. 2011). Based on these data, we proposed that transformed and immortalized cells require FACT because DNA replication in these cells is dependent on FACT. If true, then these cells would suffer from FACT loss only during DNA replication. Therefore, if the cells were arrested, they should be more tolerable of *Ssrp1* KO. To test this hypothesis, we treated immortalized and transformed cells, which were either growing or arrested (high density, low serum), with 4-OHT. Excision of *Ssrp1* (Fig. S6), did not decrease the viability of the arrested cells (Fig.3). However, when the arrested cells were plated at low density in normal serum, there was a significant reduction in the number of immortalized cells and almost no viable transformed cells within 3 to 4 days (Fig.3).

**Figure 3.**
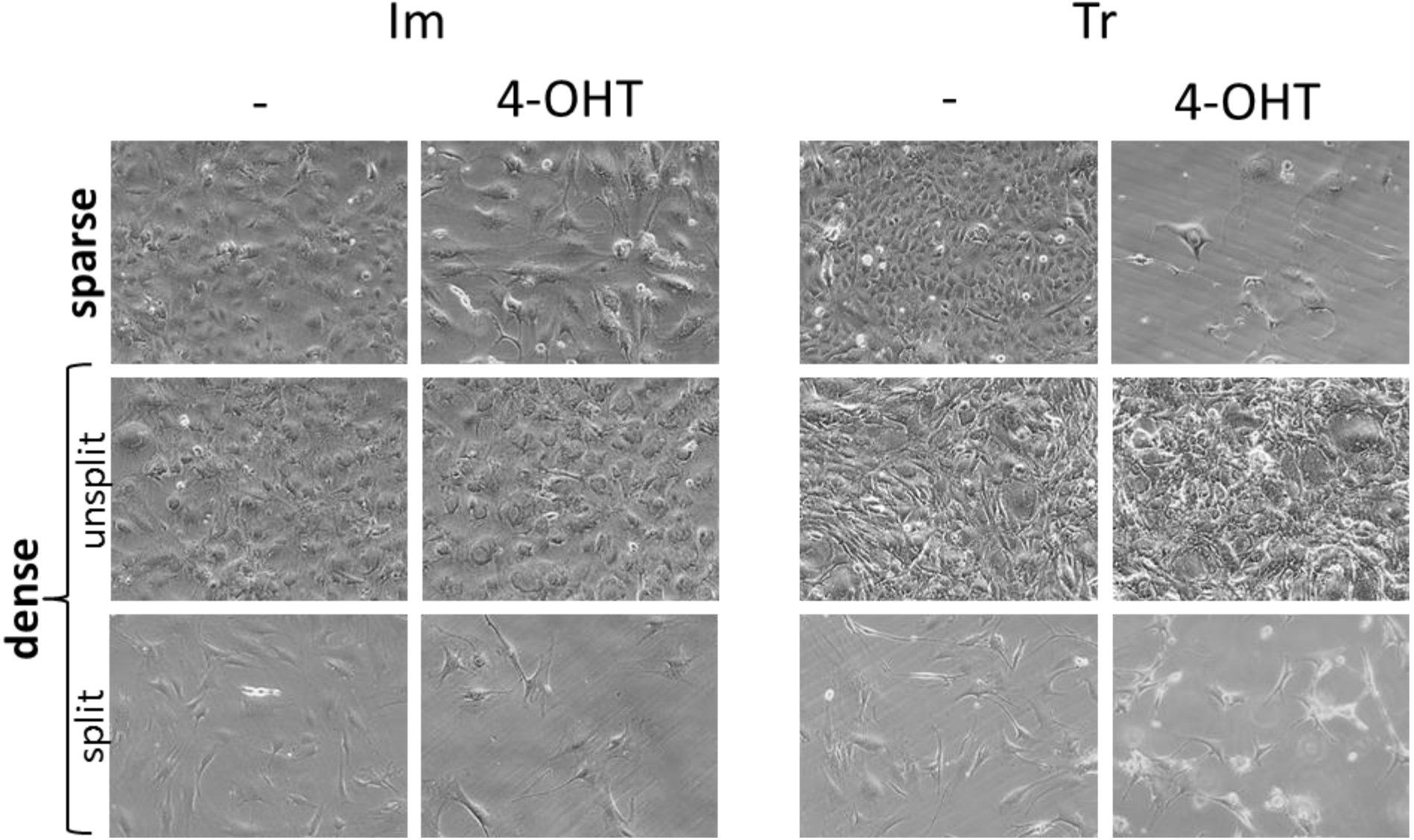
Proliferating cells are sensitive to FACT depletion. Cells were plated at two different densities: sparse (~10% confluency at the start of 4-OHT treatment) or dense (100% confluency at the start of 4-OHT treatment) and treated with 4-OHT or vehicle for five days. Representative microphotographs of Im and Tr cells taken at the end of 4-OHT or vehicle treatment for sparse cells and ten days later for dense cells. Dense cells prior to passaging were maintained in medium containing 0.5% serum.

Together, these data suggest that FACT loss results in the inability of transformed cells to pass through DNA replication, leading to DNA damage and death via apoptosis. The same trend was observed in immortalized cells, but not in primary cells. Importantly, the difference in the FACT dependence of cells cannot be explained by the difference in cell proliferation because in basal conditions, immortalized and transformed cells had similar cell cycle profiles and the same number of EdU positive cells in the populations (Fig. S3, S4).

### 3. Effect of FACT inactivation on transcription

The problem observed with replication in transformed and immortalized cells following FACT removal may be due to the direct involvement of FACT in replication or because of the loss of FACT-dependent transcription of genes involved in replication. We previously observed the enrichment of FACT on genes involved in replication and control of cell proliferation in human tumor cells (Garcia et al. 2013; Fleyshman et al. 2017).

To discriminate between these scenarios, we assessed the effect of FACT inactivation on transcription, global, using the 5-ethynyluridine (EU) incorporation assay, and of individual genes using next-generation sequencing (NGS) of RNA isolated from cells (RNA-seq). Surprisingly, the EU incorporation was increased following *Ssrp1* KO in immortalized and transformed cells but not changed in primary cells. No changes in EU incorporation were observed in any of the MSF cell types from mice heterozygous for *Ssrp1^fl/+^* alleles (Fig. 4A and S7).

**Figure 4.**
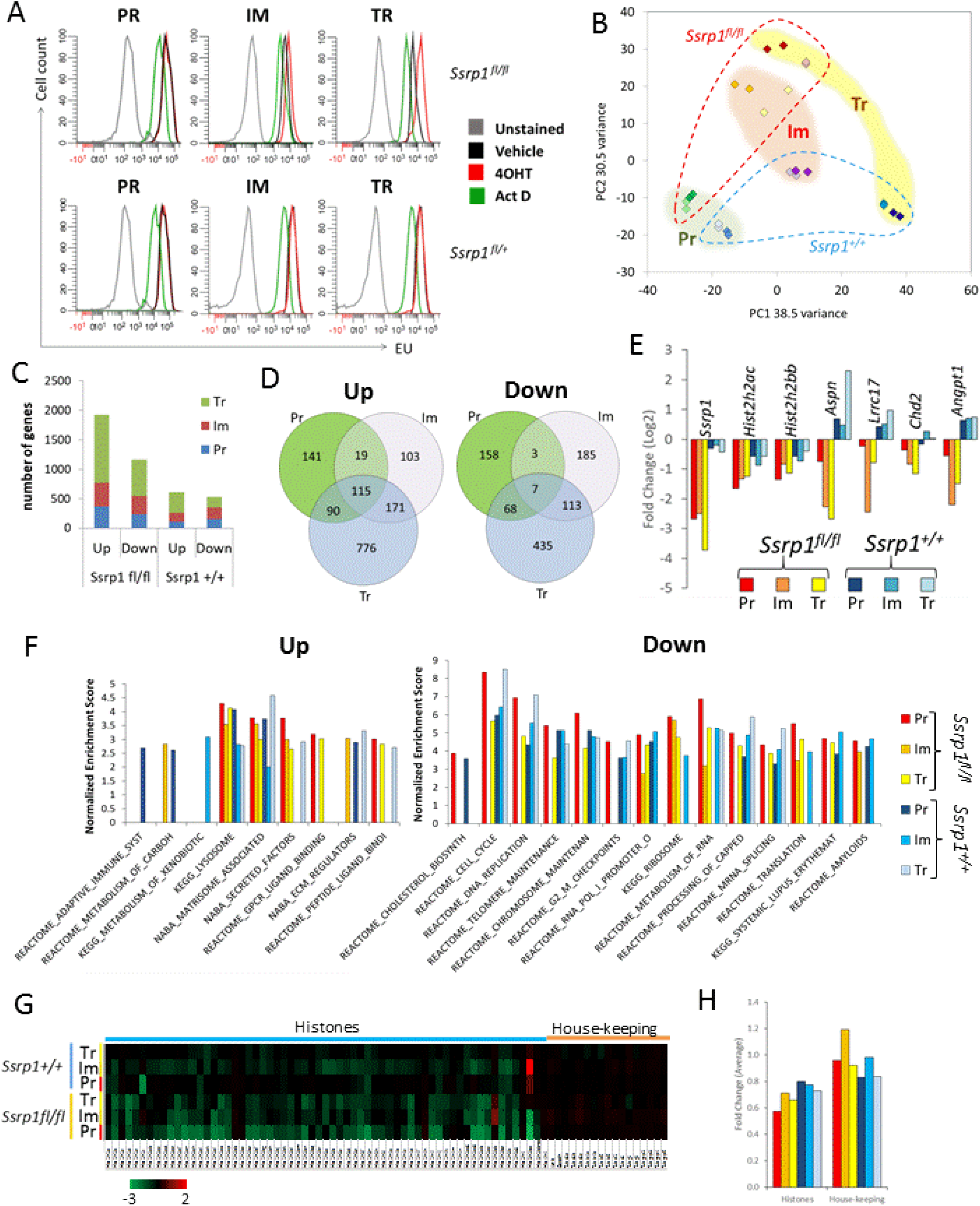
Effect of FACT depletion on transcription in mouse cells. A. EU incorporation into primary (Pr), immortalized (Im), and transformed cells (Tr) of two genotypes treated with 4-OHT or vehicle for five days. Un stained cells and cells treated with the transcription inhibitor Actinomycin D (ActD) were used as controls. Data for one representative cell line of each genotype are shown. Data for other cell lines are shown in Fig. S6. B-H. Analyses of two biological replicates of paired-end RNA-seq samples from Pr, Im and Tr homozygous wild-type or floxed *Ssrp1* cells treated with 4-OHT or vehicle. The correlations between replicates are shown on Fig. S8. B. Principle component analysis. Darker colors, 4-OHT; lighter colors, control samples. C. Number of upregulated and downregulated genes. Fold change (FC) > ±1.5, adjusted p-value < 0.05. D. Differentially upregulated and downregulated genes shared by Pr, Im, and Tr cells with floxed *Ssrp1* following 4-OHT treatment. Venn diagrams for wild-type Ssrp1 cells are shown on Figure S8. E. Fold change in the expression of the genes downregulated in all three cell types with floxed *Ssrp1* following 4-OHT treatment. F. Summary of GSEA. Normalized enrichment scores (NES) are shown for all significantly enriched gene lists in Pr, Im or Tr cells of either of genotype. Individual NES plots are shown on Fig. S9. G. Heat plot demonstrating the FC in the expression of replication-dependent histones and several housekeeping genes in different conditions in cells treated with 4-OHT or vehicle. H. Average FC in expression of all replication-dependent histones and selected housekeeping genes presented in panel G for cells treated with 4-OHT or vehicle.

For RNA-seq, we used cells from *Ssrp1^fl/fl^; CreER^T2+/+^* and *Ssrp1^+/+^; CreER^T2+/+^* mice to filter out the effect of 4-OHT administration and Cre activation, independent of Ssrp1 KO. Cells for RNA isolation were collected after five days of 4-OHT treatment before any visible signs of toxicity appeared in transformed and immortalized cells from the *Ssrp1^fl/fl^; CreER^T2+/+^* mice. Two biological replicates were used for each condition, and their correlation was >98% (>99% in 11 out of 12 cases, Fig. S8). Principal component analysis (PCA) confirmed the similarity of all replicates (Fig. 4B). In addition, PCA showed that all primary samples were clustered together independently of their *Ssrp1* alleles. The administration of 4-OHT had little effect on the primary cells of either genotype (Fig. 4B). The immortalized and transformed samples diverged from the primary MSFs in two directions: samples from cells with wild-type *Ssrp1* were found along PC1 (38.5% variance); and samples from cells with floxed *Ssrp1* alleles were found along PC2 (30.5% variance). The 4-OHT-treated samples from immortalized and transformed cells with floxed *Ssrp1* were clearly separated from the untreated samples, but the impact of 4-OHT administration on the distribution of these samples was very modest and shifted the immortalized and transformed samples closer to the position of the primary samples along PC1 (Fig. 4B). The major difference between the transcriptional profiles of all tested cells appeared to be due to immortalization and transformation. Not surprisingly, the latter (p53 inactivation plus mutant HRas^V12^ overexpression) had a stronger effect than p53 inactivation alone. Thus, PCA showed that transcription of the cells was changed much stronger due to immortalization and transformation, than due to *Ssrp1* KO. The changes were of a rather random nature as there was a high variance between independently generated immortalized and transformed cells. Because the differences were observed even before *Ssrp1* deletion, it cannot be attributed to FACT inactivation. In line with the phenotypic studies, FACT inactivation had a more prominent effect on gene expression in transformed and immortalized cells than in primary cells.

Because Cre is known to cause random DNA breaks and a DNA damage response (Loonstra et al. 2001; Janbandhu et al. 2014), and 4-OHT binds and inhibits the estrogen receptor (Quirke 2017), the identification of FACT-dependent genes should be performed by comparing the gene expression profiles of cells with floxed and wild-type *Ssrp1* alleles that both harbor CreER^T2^ and were similarly treated with 4-OHT. However, taking into account the difference between cells with wild-type and floxed *Ssrp1* even before 4-OHT treatment using PCA (Fig. 4B), we chose to first compare the same cells of both genotypes with and without 4-OHT treatment and then compare the gene expression changes between *Ssrp1*-floxed and wild-type cells to filter out the effects of 4-OHT administration and Cre activation.

After 4-OHT administration, there were more upregulated than downregulated genes in all cell types with floxed *Ssrp1*. In contrast, cells with wild-type *Ssrp1* had similar numbers of upregulated and downregulated genes (Fig. 4C). There was little overlap between the gene expression changes observed with 4-OHT administration of cells with floxed and wild-type *Ssrp1* except for the genes downregulated in the primary cells (Fig. S9). There were 115 genes upregulated across all three cell types following *Ssrp1* KO but only seven that were downregulated (Fig. 4D). The most significantly and specifically (i.e., only in *Ssrp1*-floxed but not wild-type cells) downregulated gene was Ssrp1 itself (Fig. 4E). The commonly upregulated or downregulated genes did not belong to any known gene sets based on gene set enrichment analysis (GSEA).

Surprisingly, when we performed GSEA on all upregulated or downregulated genes in cells with wild-type and floxed *Ssrp1* after 4-OHT administration, we obtained very similar lists for all cell types (Fig. 4F and S10), suggesting that 4-OHT administration and Cre activation, but not *Ssrp1* KO, might be the major driver of the gene expression changes. No functional gene categories specifically responded to *Ssrp1* KO in the cells in which we observed strong phenotypic changes (i.e., immortalized and transformed cells with floxed *Ssrp1*). There was significant enrichment of genes involved in cell cycle-related processes (e.g., DNA replication, telomeres, and chromosome maintenance) among genes downregulated by 4-OHT. However, their expression was reduced in both *Ssrp1* floxed and wild-type cells. Downregulation of these genes was stronger in cells with *Ssrp1* floxed than wild-type alleles. For example, expression of the replication-dependent histone genes, which are highly expressed during the S-phase of the cell cycle, was reduced to a greater extent in *Ssrp1* floxed cells than *Ssrp1* wild-type cells (Fig. 4F and G), which is consistent with the observed phenotypic changes. However, the strongest reduction was observed in primary cells in which we saw no difference in growth or cell cycle distribution upon 4-OHT treatment. Because FACT inactivation caused an enrichment of the same gene lists among the upregulated and downregulated genes, FACT loss may result in alterations of the same process or pathway. Alternatively, the low number of genes for which expression was similarly changed in the different cell types suggests that changes in gene expression may be secondary to FACT loss (i.e., in response to dysregulation of certain processes rather than direct FACT involvement in the transcription of these genes).

To test this hypothesis, we investigated whether FACT was present at genomic regions of genes for which expression changed upon *Ssrp1* KO and whether the transcription of genes that were enriched for FACT was changed upon FACT loss. We performed chromatin immunoprecipitation with an SSRP1 antibody followed by NGS (ChIP-seq) in immortalized and transformed cells. The level of FACT in the primary cells was too low to accurately run in this assay. We observed SSRP1 enrichment at coding regions of genes in proportion to the level of transcription of these genes (Fig. 5A). In immortalized cells, this dependence was much more evident than for transformed cells, where there was more SSRP1 at non-transcribed regions and genes expressed at low levels. Therefore, the difference in SSRP1 enrichment depending on transcription was less pronounced (Fig. 5A). This pattern was reproduced with independently generated immortalized and transformed cells from a different mouse and using a different platform for RNA-seq (Fig. S11, see details in Material and Methods). To confirm these findings, we assessed the correlation between SSRP1 and RNA read coverage under basal conditions and observed a stronger positive correlation with immortalized cells compared to the transformed cells (Fig. 5B and S12A). However, the distribution of nucleosomes measured using histone H3 ChIP-seq was not different between the two cell types (Fig. S11).

**Figure 5.**
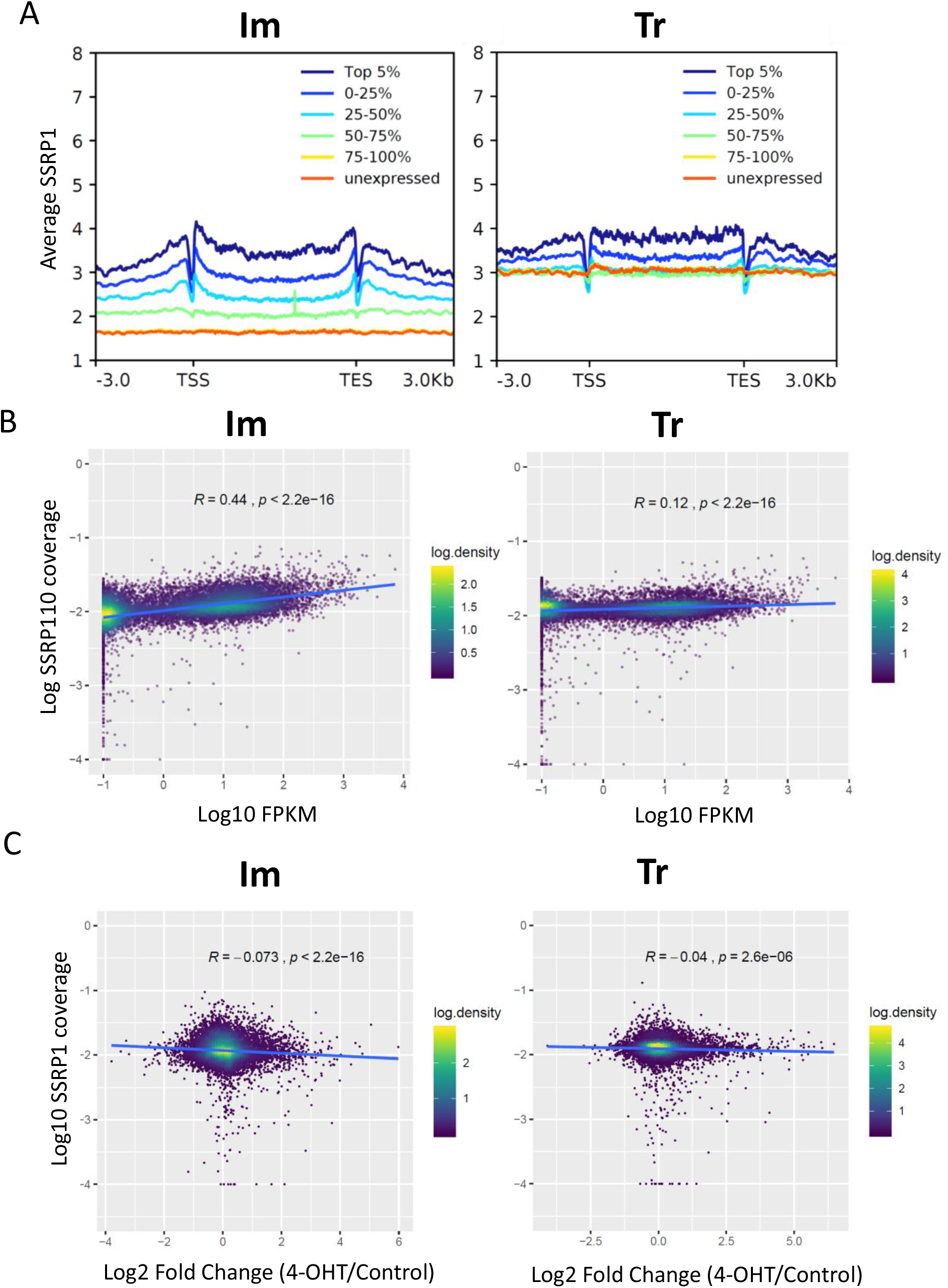
FACT-dependence of gene transcription in mouse immortalized (Im) and transformed (Tr) cells. Data are presented for *Ssrp1^fl/fl^; CreER^T2+/+^* #2. Data for other cell lines are shown on Figs. S11 and S12. A. Average enrichment of SSRP1 (ChIP-seq) at genes depending on the levels of their transcription defined by RNA-seq read density in basal conditions. B. Dot plot demonstrating the relationship between SSRP1 enrichment and transcription. C. Dot plot demonstrating the relationship between SSRP1 enrichment and fold change in gene expression following *Ssrp1* KO. R – Pearson correlation coefficient.

Next, we compared the FACT-dependence of the transcription of genes occupied by FACT by correlation analysis between FACT coverage based on ChIP-seq data and fold change in gene expression following *Ssrp1* KO. In all cases, we saw a weak negative correlation between FACT enrichment and the FACT-dependence of gene expression (Fig. 5C and S12B, C). Although weak, these correlations were highly significant and stronger for immortalized cells than for transformed cells (i.e., loss of FACT led to a stronger increase in gene expression in immortalized cells than in transformed cells), suggesting that the presence of FACT might interfere with gene transcription, and this effect is weakened in transformed cells.

### 4. Effect of FACT inactivation on transcription in human cells

While preparing this manuscript, two papers were published showing that FACT inactivation did not lead to the repression of the expression of genes enriched for FACT in mouse embryonic stem cells and normal human fibroblasts (Kolundzic et al. 2018; Mylonas and Tessarz 2018). Therefore, we examined the relationship between FACT enrichment and transcription in human tumor cells using the fibrosarcoma cell line HT1080 for which we already had SSRP1 ChIP-seq data from two independent experiments of 2 to 3 replicates (Garcia et al. 2013; Nesher et al. 2018) and data from several gene expression studies using different methods for FACT knockdown (KD) (e.g., SSRP1 shRNA, siRNAs to SSRP1 and SPT16). We also had nascent RNA-seq data from the same cells (Nesher et al. 2018), which served as a more accurate measure of gene expression that was not influenced by the degree of RNA stability (Fig. S13). As we previously observed, there was a significant positive correlation between FACT enrichment and the transcription of genes under basal conditions in these cells (Fig. 6A). Interestingly, this dependence was not only quantitative but also qualitative because the profiles of FACT enrichment were different for genes expressed at different levels (Fig. 6B). The top 5% of the expressed genes had the highest level of FACT at the coding regions, and two peaks at the 5’-UTR and 3’-UTR separated by FACT depleted regions at transcription start and end sites (TSS and TES), which is in line with nucleosome depletion at these regions (Fig. 6B). The top 25% of the expressed genes had much less FACT at the coding regions than at the 5’UTR. The coding regions of this group of genes had FACT present only just after the TSS. Other expressed genes had FACT almost exclusively at the 5’UTR followed by a sharp reduction of FACT immediately after the TSS. There was some enrichment of FACT at the coding regions of untranscribed genes (Fig. 6B).

**Figure 6.**
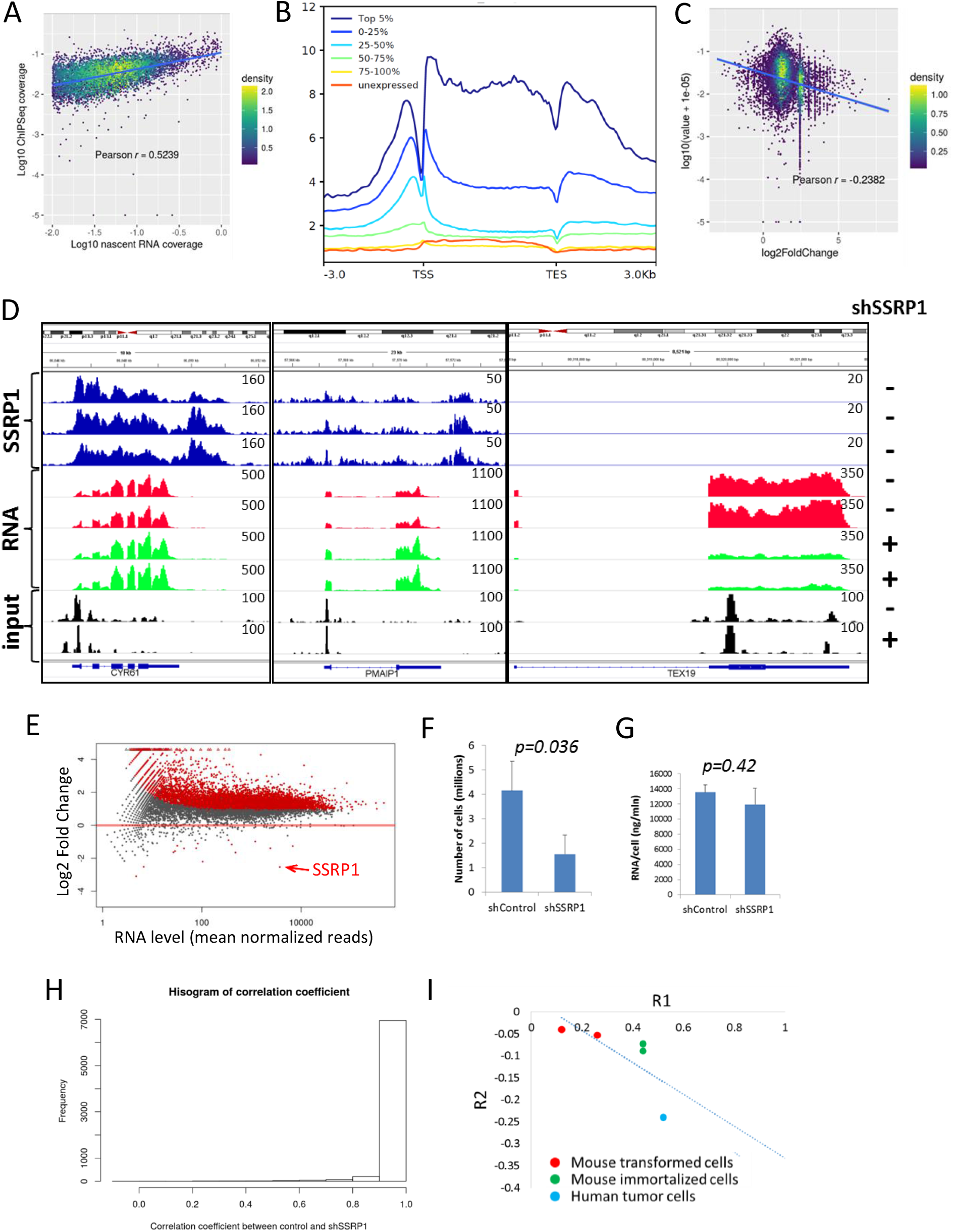
Dependence of gene transcription on the presence of FACT on chromatin in HT1080 human fibrosarcoma cells. A. Dot plot demonstrating the relationship between SSRP1 enrichment (ChIP-seq) and transcription in basal conditions (nascent RNA-seq). R – Pearson correlation coefficient. B. Average enrichment of SSRP1 (ChIP-seq) at genes depending on the levels of their transcription defined by nascent RNA-seq read density in basal conditions. C. Dot plot demonstrating the relationship between SSRP1 enrichment and fold-change in gene expression (RNA-seq) upon SSRP1 knockdown with shRNA. D. Integrated genomic view of normalized read density of SSRP1 (ChIP-seq, three replicates), RNA (RNA-seq, two replicates), and input DNA (two replicates) in cells transduced with control shRNA (-) or shRNA to SSRP1 (+) at three genomic regions surrounding *CYR61, PMAIP*, and *TEX19*. E. Changes in gene expression between shSSRP1 and shControl cells depending on the level of expression in basal conditions using loess normalized counts. Red dots – fold change > ±1.5, adjusted p-value <0.05. F. Number of cells transduced with control or SSRP1 shRNA 72 h after transduction. The bars represent the mean ± SD (n = 2). G. The amount of RNA isolated from cells transduced with control or SSRP1 shRNA 72 h after transduction. The bars represent the mean ± SD (n = 2). Data for the individual replicates are shown in Fig. S15. H. Comparison of RNA-seq reads corresponding to the same exons between shSSRP1 and shControl samples. I. Inverse relationship between two correlation coefficients, R1, SSRP1 enrichment and gene transcription in basal conditions, and R2, SSRP1 enrichment and change in gene expression upon FACT depletion, for all tested cells.

We performed correlation analysis between FACT enrichment and changes in gene expression upon FACT KD. As before, we saw only a negative correlation independent of what data was used (i.e., microarray hybridization or RNA sequencing), the type of KD (sh or siRNAs to SSRP1, SUPT16H, or both), or whether the correlation analysis was performed using SSRP1 enrichment only at the promoter or coding regions or both (Fig. 6C and S14, S15). For the RNA-seq, we used spike-in controls that allowed us to detect absolute changes in transcription. Consistent with the mouse data, FACT downregulation led to an increase in the transcription of multiple genes. There were more genes enriched for FACT among the upregulated genes than the downregulated genes (Fig. 6D, E). The negative correlation between FACT enrichment and changes in gene expression upon FACT KD was much stronger in these cells than in mouse (Fig. 6F). We also observed that although the number of HT1080 cells was significantly reduced upon FACT KD, cells without FACT had the same amount of RNA per cell as the control cells (Fig. 6G, H and S16). Together, these data confirmed that in human tumor cells, loss of FACT is not accompanied by a reduction in transcription.

Cell-free experiments showed that the absence of FACT caused RNA polymerase II to pause at several positions within the nucleosome and, therefore, produce early terminated short transcripts instead of fully functional transcripts (Hsieh et al. 2013). In addition, several studies in yeast demonstrated that inhibition of FACT was associated with the loss of nucleosomes at the coding regions of genes, which might lead to cryptic intra-genic initiation (Morillo-Huesca et al. 2010; Myers et al. 2011; Erkina and Erkine 2015; Feng et al. 2016). In both cases, non-functional transcripts (i.e., early terminated or incorrectly initiated) might mask the reduced presence of proper full-length transcripts when analyzed using NGS (short reads) or total EU incorporation. However, both types of shortened transcripts could skew the distribution of reads along the coding region: early termination could generate bias towards an overrepresentation of reads corresponding to the 5’ versus 3’ exons; cryptic initiation could result in an overrepresentation of reads from the 3’ exons. Therefore, we looked for a correlation between the number of reads representing individual exons between samples in the presence or absence of FACT, and the data was not skewed (Fig. 6H).

For the limited number of cases tested, we observed an inverse relationship between the positive correlation coefficient for FACT enrichment at genes versus the transcription in basal conditions and the negative correlation coefficient for FACT enrichment versus the change in expression of genes upon FACT inactivation (Fig. 6I). These data suggest that the influence of FACT on the transcription of genes is dependent on the type of cell. In general, the stronger was the FACT association with transcribed genes, the higher was the activation of transcription upon FACT inactivation in cells.

### 5. FACT inactivation leads to destabilization of chromatin in immortalized and transformed cells

The ability of FACT to bind to the nucleosome components provides the basis for our model in which nucleosome/chromatin structure may be preserved at transcribed regions by FACT. This model implies that the loss of FACT leads to the destabilization of chromatin and suggests a mechanism by which the loss of FACT may lead to increased transcription. Namely, the disassembly of nucleosomes caused by RNA polymerase passage should make passage of the next RNA polymerases easier. We used two approaches to test this hypothesis. First, we compared the sensitivity of chromatin to nuclease digestion in cells in the presence or absence of FACT. After 4-OHT treatment, both immortalized and transformed cells with *Ssrp1* floxed alleles were more sensitive to digestion with micrococcal nuclease (MNase), which preferentially cleaves protein-free DNA. There was an increase in the proportion of lower molecular weight DNA fragments upon digestion and a reduction in the length of the DNA fragments protected by the nucleosomes, indicating nucleosome opening (Fig. 7A-C and S17). No any of these changes were seen in primary cells (Fig.S17, D, E).

**Figure 7.**
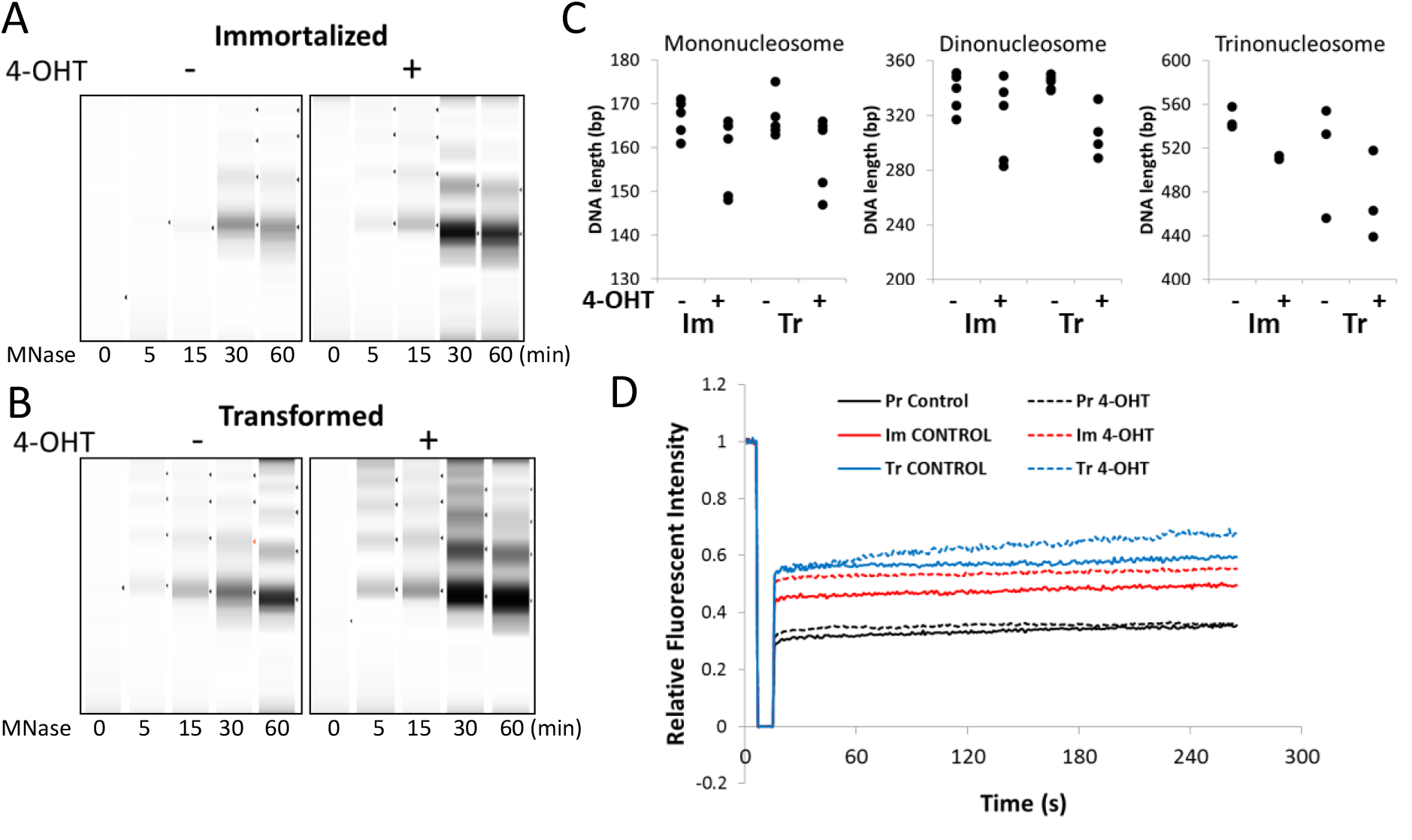
Changes in chromatin stability and dynamics following FACT depletion in mouse cells. A. MNase sensitivity of chromatin from Im and Tr cells containing floxed Ssrp1 before and after 4-OHT treatment. Represented images of capillary electrophoresis generated by the Bioanalyzer. Data are presented for one Im and one Tr cell line. Data for other cells lines are shown on Fig. S16. C. Distribution of fragments corresponding to mono-, di-, and tri-nucleosome lengths obtained following MNase digestion of chromatin from 4-OHT-treated and control Im and Tr cells detected by the Bioanalyzer. Dots represent data obtained from several independent experiments with cells isolated from different mice. D. Mean relative fluorescent intensity before and after photobleaching of Pr, Im, and Tr cells treated with 4-OHT or vehicle. Curves represent the mean of 15 to 20 curves obtained from individual cells and normalized to the intensity of the pre-bleach fluorescence (five measurements). Post-bleach fluorescence was measured every second during 250 seconds. The standard deviation between cells from the same sample were below 10% and omitted for clarity. Data for other cells of both genotypes are shown on Fig. S17.

For the second approach, we transduced cells with lentivirus encoding mCherry-tagged histone H2B and then compared cells treated with 4-OHT with untreated control cells using the fluorescence recovery after photobleaching (FRAP) assay to measure the dynamics of histone turnover within the chromatin. The recovery of the fluorescence at the site of bleaching occurs due to the exchange of bleached histones for histones from unbleached nuclear regions. Almost immediate recovery occurs due to the incorporation of free histones present in the nucleoplasm, and slow recovery is due to the eviction of histones from unbleached chromatin. All experiments were done within 300 seconds. Therefore, new histone synthesis was considered neglectable. Consistent with the MNase data, there was a fast recovery observed in immortalized and transformed cells containing the floxed *Ssrp1* allele upon *Ssrp1* KO, which did not occur in cells with wild-type *Ssrp1*, suggesting a larger pool of free histones and higher histone mobility in the absence of functional FACT (Fig. 7D and S18).

Both approaches demonstrated that in immortalized and transformed mammalian cells, FACT is critical for the stabilization of chromatin and prevention of histone loss from chromatin.

### 6. Why FACT is critical for viability of transformed, but not of primary cells

Many of the changes in transformed cells resulting from FACT KO were not as prominent or absent in primary cells. Immortalized cells demonstrated intermediate changes with some cell lines acting more primary cell-like and others transformed cell-like. These data suggest that either FACT performs a specific function in transformed cells or its function is underutilized in primary cells compared to transformed cells. The two most prominent effects of FACT loss that were observed almost exclusively in transformed cells were the impairment of DNA replication (reduced EdU incorporation, G2/M increase, and appearance of anaphase bridges) and chromatin destabilization (increased sensitivity to MNase, elevated histone dynamic).

All cells used in the study proliferated albeit at different rates. Moreover, different lines of immortalized and transformed cells proliferated at different rates. However, FACT loss was still more toxic for transformed cells compared to immortalized cells and non-toxic for proliferating primary cells, suggesting that replication was not the major condition of cell dependence on FACT. To determine what else could make cells sensitive to FACT loss, we compared the stability of chromatin between primary, immortalized, and transformed cells. FRAP experiments clearly showed that the basal level of histone mobility was higher in transformed than immortalized cells, which was higher than in primary cells. These observations were made in cells with both floxed and wild-type *Ssrp1* (Fig. 7D and S18). Transformed cells were also more sensitive to MNase digestion than immortalized cells (Fig. 7A, B and S17).

From the analyses of the RNA-seq data, we observed significant changes in the proportion of reads with assigned genomic features versus no features between primary, immortalized, and transformed cells independently of genotype and 4-OHT treatment (Fig. 8A,B). There were significantly more reads with no features (i.e., genomic regions lacking any known transcripts or regulatory elements) in immortalized and transformed cells compared to the primary cells. The appearance of these reads may suggest either contamination of RNA with genomic DNA or that the samples contained elevated levels of products of so-called pervasive or illicit transcription from non-coding genomic regions due to the loss of chromatin packaging at these regions (Layer and Weil 2009). Because the RNA was isolated simultaneously using the same method for all types of cells, the first explanation (i.e., genomic DNA contamination) seems less probable. However, to exclude this possibility, we analyzed the samples from independently isolated, immortalized, and transformed cells from mice with floxed *Ssrp1* using a different method of RNA isolation (Trizol reagent versus purification column) and sequencing platform (single versus paired-end reads). The data generated was consistent with the first set of data (Fig.8 C,D).

**Figure 8.**
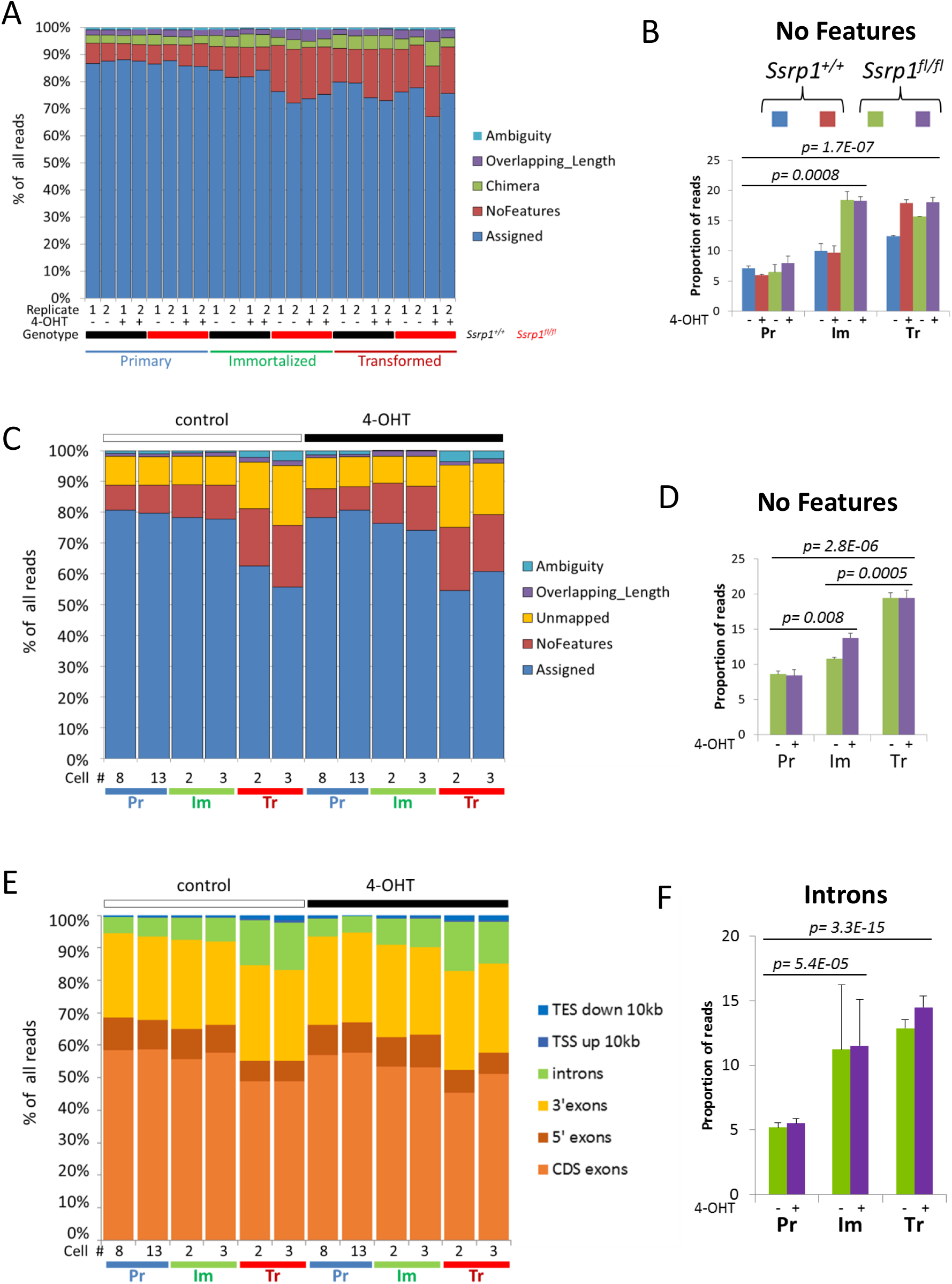
Increased pervasive transcription in cells following immortalization and transformation. A-D. Distribution of RNA-seq reads corresponding to annotated genomic features (Assigned) or lack of features (NoFeatures). and reads with questionable annotations (Ambiguity, Chimera, Overlapping_Length) for RNA-seq samples from two independent experiments. A-B. RNA from Pr, Im, and Tr cells of two genotypes treated with 4-OHT or vehicle was prepared using a column-based method of RNA isolation and sequenced as 100 bp paired-end reads. A. Distribution of reads within individual samples. B. Mean percentage of reads with no features between replicate samples +/-SD (n=2). C-D. RNA from Pr, Im, and Tr cells with floxed Ssrp1 treated with 4-OHT or vehicle prepared using a TRIZOL-based RNA isolation method and sequenced as 75 bp single-end reads. C. Distribution of reads within individual samples. D. Mean percentage of reads with no features ± SD (n = 2). E-F. Distribution of single-end RNA-seq reads corresponding to genic features. E. Distribution of reads in individual samples. F. Mean percentage of reads corresponding to introns ± SD (n = 2). Data for paired-end reads are shown on Fig. S19.

Furthermore, in both RNA-seq datasets, we observed a significant increase in the proportion of reads corresponding to introns in transformed and, to a lesser degree, immortalized cells (Fig. 8C and S19). This so called “intron retention” was previously observed in tumor cells (Dvinge and Bradley 2015; Smart et al. 2018) and was thought to be the result of mutations in splicing factors (Wong et al. 2016). However, it can also be explained by cryptic initiation from introns due to the loss of nucleosomes. Because we detected “intron retention” in all independently transformed cell cultures concomitant with other signs of chromatin destabilization, we propose that the second explanation is more probable.

Our data suggest that upon immortalization and transformation, chromatin in cells becomes less condensed based on higher sensitivity to nuclease digestion, more dynamic based on an enlarged pool of free histones, and more frequent histone exchange between different nuclear regions, demonstrated by FRAP, and more accessible for transcription machinery based on the elevated appearance of the products of pervasive transcription. All these effects were much stronger in transformed cells compared to immortalized cells, suggesting a general loss of chromatin stability upon transformation.

## Discussion

The goal of our study was to understand why malignant tumor cells are much more dependent on FACT than less malignant and non-tumor cells. The problem that FACT function established in cell-free experiments, facilitation of RNA polymerase passage though chromatin, makes difficult to explain differential cell dependence on FACT, since nucleosomes are present at coding regions of all genes in all mammalian cells. Thus we designed this study to compare consequences of FACT loss in original FACT-independent cells and cells which we made dependent on FACT via established genetic manipulations, i.e. inactivation of tumor suppressor p53 and overexpression of mutant HRas^V12^ oncogene. These cells as expected became fully transformed, i.e. anchorage independent and forming aggressive tumors in mice, thus representing a model of malignant tumor cells.

Our initial hypothesis was that FACT is needed not for all transcription elongation, but for the most efficient high-rate transcription. There were data in literature that transcription is elevated upon Ras-induced transformation (Kotsantis et al. 2016). Indeed we observed increase in general transcription upon immortalization and transformation of the cells in our study. In support of this hypothesis, we and others observed correlation between FACT enrichment genome-wide and rates of transcription (Chang et al. 2018; Kolundzic et al. 2018; Mylonas and Tessarz 2018). However, inactivation of FACT did not reduce rates of transcription in any of tested cells, ruling out proposition that FACT is needed for general transcription elongation.

Although there were some genes with reduced expression following loss of FACT, these genes were not located in genomic regions that were enriched with FACT under basal conditions. Moreover, there was a negative correlation between FACT enrichment and changes in gene transcription, suggesting that FACT inhibited the transcription of these genes.

Thus FACT enrichment genome-wide depends on transcription, but transcription does not depend on FACT, suggesting that FACT function at transcribed regions is different from transcription elongation. Indeed, our studies and the existing FACT literature suggest that instead of disassembly, FACT preserves the nucleosomes that are disturbed by the passage of the RNA polymerase (Gurova et al. 2018). If RNA polymerase itself or together with other factors destabilizes nucleosomes, the stabilization or reassembly of nucleosomes by FACT will prevent the loss of histones and their associated epigenetic information. Furthermore, the model explains why FACT is enriched in proportion to the level of transcription. As more RNA polymerases pass through a gene, the more the nucleosomes are disturbed, which uncovers more FACT binding epitopes.

Our current model of FACT function is consistent with published structural studies that showed that mammalian FACT could only bind nucleosomal components or partially disassembled nucleosomes (e.g., lacking the H2A/H2B dimer) because the FACT binding epitopes are hidden within the folded nucleosome (Tsunaka et al. 2016; Safina et al. 2017; Wang et al. 2018). It is also consistent with studies in yeast that showed that FACT is targeted to transcribed chromatin through its recognition of RNA polymerase-disrupted nucleosomes (Martin et al. 2018) and the loss of FACT is accompanied by the loss of histones and pervasive transcription (Morillo-Huesca et al. 2010; Myers et al. 2011; Erkina and Erkine 2015; Feng et al. 2016). Similar observations in plants have also been recently reported (Nielsen et al. 2018).

If the above model is correct, why is transcription increased following the loss of FACT? There are two potential explanations for this phenomenon, which will require further investigation. First, in the absence of FACT, nucleosomes that are destabilized by the first RNA polymerase passage will be a weaker barrier for subsequent passages by RNA polymerases. Thus, gene transcription will become more efficient. Alternatively or in addition, nucleosomes may eventually be lost from the transcribed regions, leading to cryptic initiation and pervasive transcription. With short read-based transcription analysis, it is difficult to distinguish between these two scenarios. We expected that cryptic initiation and pervasive transcription would skew the distribution of RNA-seq reads corresponding to the 5’ and 3’ exons of the genes. However, we observed a very high correlation between the exon reads in the control and FACT-depleted cells, which suggests more efficient transcription as a short-term effect of FACT loss.

FACT loss was more toxic to proliferating cells than quiescent transformed cells, suggests that FACT is involved in replication as was previously noticed (Tan et al. 2006; Tan et al. 2010; Abe et al. 2011). However, most published studies concluded that FACT disassembles nucleosomes in front of the replisome. If this is the case, FACT has almost opposite effects on replication and transcription. If FACT reassembles nucleosomes during replication as it does during transcription, then why would replication be decreased in FACT-depleted cells? One proposed reason is that the replisome slows down if nucleosomes are not correctly or timely assembled behind it; however, the mechanism is currently unclear (Groth et al. 2007). In a separate study we present data and propose explanation of why FACT is required for replication in transformed cells (Prendergast L. et al, in preparation).

The most interesting question is why FACT loss is only problematic for transformed and tumor cells if it serves a very basic function of prevention the loss of histones from destabilized nucleosomes during transcription and replication. Although both processes occur in primary cells, FACT loss is not associated with reduced viability of these cells and destabilization of chromatin. One explanation is that replication and transcription occur at a lower rate in the primary cells than in the immortalized and transformed cells, and therefore the rate of these processes could be one factor that determined the necessity of FACT. However, this hypothesis was disproven by a recent report that demonstrated increased proliferation of mouse ESCs, which typically have very high replication and transcription rates (Efroni et al. 2008), upon FACT knockdown (Mylonas and Tessarz 2018).

Another hypothesis of why FACT is essential for transformed and tumor cells is that it is needed for the packaging of DNA for mitosis, which is a constantly ongoing process in these cells. FACT was found to be one of only a few factors that were essential for packaging mitotic chromosomes under cell-free conditions (Shintomi et al. 2015). However, the ability of primary and other non-tumor cells to pass through mitosis in the absence of FACT shakes this proposition.

The third hypothesis is that FACT prevents nucleosome loss in transformed and tumor cells, in which chromatin is already destabilized comparing with non-tumor cells. We showed that both transformation and FACT loss reduced nucleosome stability. Nucleosome stability is a well-defined property of nucleosomes in cell-free experiments, which can be measured by different methods, including resistance to increased concentrations of salt and protection of nucleosomal DNA from nuclease digestion. However, in mammalian cells, there are more than a million nucleosomes, and their stability differs significantly at different genomic loci and depends on a large number of factors. Therefore, understanding nucleosome stability in the context of cells is more difficult. To make this concept easier, we propose to define nucleosome stability in cells as a degree of this nucleosome interference with transcription. In general, regions of constitutive heterochromatin in mammalian cells have more stable nucleosomes (Collings et al. 2013; Riedmann and Fondufe-Mittendorf 2016) whereas nucleosomes in transcribed regions are less stable and more open (Brower-Toland et al. 2005). Nucleosomes at AT-rich DNA (e.g., promoters and TSS regions) are also less stable (Lorch et al. 2014).

Based on indirect literature data and observations made in the current study, we propose that, in general, chromatin is destabilized upon malignant transformation. It becomes more sensitive to nucleases, has a higher histone exchange rate, and is less restrictive to transcription (judged by the appearance of reads with no features and corresponding to introns). Data available in the literature suggest that changes in the chromatin state may be a universal process that accompanies malignant transformation. Reports have shown that tumors can have reduced expression of linker histone 1 (Scaffidi 2016), hypomethylated DNA (Ehrlich 2009), changes in the expression of architectural chromatin proteins (e.g., reduction of HP1) that could make chromatin less stable, (Dialynas et al. 2008), and increased levels of HMG box proteins (Hock et al. 2007). In addition, we observed a higher sensitivity of tumor cells to chromatin-destabilizing small molecules, which suggests that chromatin in tumor cells may be less stable under basal conditions than in normal cells (Gasparian et al. 2011; Safina et al. 2017).

The reasons for chromatin destabilization in tumor cells are not clear, but some can be easily proposed, such as elevated rates of replication and transcription. Conversely, the elevated rates of transcription and replication in tumor cells may be possible due to less stable chromatin. Moreover, there may be significant benefits of less stable chromatin in tumors because it may permit easier changes of transcriptional programs in tumor cells, which could account for the phenotypic plasticity and easiness of transitioning between epigenetic states as observed in malignant tumor cells.

We observed an interesting relationship between FACT and transcription in immortalized and transformed cells, which was different from that observed in normal mammalian stem and non-stem cells (Kolundzic et al. 2018; Mylonas and Tessarz 2018). In immortalized cells, FACT enrichment was more proportional to the level of gene transcription than that observed in transformed cells. When FACT was depleted from the immortalized cells, there was a higher increase in transcription of the FACT-enriched genes compared to the transformed cells. Based on these data, the ability of FACT to rebuild nucleosomes at transcribed genes may be stronger in immortalized cells than in transformed cells. In transformed cells, the levels of FACT and FACT enrichment at non-transcribed genes and non-genic regions are higher. Because chromatin is less stable in transformed cells, there is broader pervasive transcription. Thus, FACT function is needed not only at transcribed genes, but genome-wide, which results in FACT enrichment genome-wide.

On the background of already destabilized chromatin, the loss of FACT may further destabilize the chromatin, leading to fatal consequences for transformed and tumor cells. Currently, however, we know very little about the consequences of chromatin destabilization in cells. We recently proposed several mechanisms (Gurova, BioEssays, 2019, accepted) but they have not been explored experimentally.

Taken together, our findings suggest that FACT is essential for tumor cells to compensate for the general destabilization of chromatin, which cells acquire during the process of transformation.

## Materials and Methods

### Materials and reagents used in the study are listed in Table S1

#### Phenotypic characterization of consequences of *Ssrp1* KO in mouse cells

MSFs (1-2 × 10^6^ cells) were plated in 150 mm plates. The next day, cells were treated with 2 μM 4-OHT for 96 h (Pr) or 120 h (Im and Tr). The medium was replaced every 48 h with fresh 2 μM 4-OHT. At the end of treatment, both treated and untreated cells were trypsinized and re-plated for further experiments. *Ssrp1* excision was confirmed using western blotting or genomic PCR as described (Sandlesh et al. 2018). Experiments were performed in triplicate and repeated at least twice.

Cell viability was assessed five days after plating of 5 × 10^5^ per well of a 6-well plate using Cell Titer Blue Reagent (Promega). Cell cycle analysis was performed as previously described (Gasparian et al. 2011). For 3D colony growth, 1 × 10^5^ cells were mixed with 0.3% agarose and plated over a layer of 0.5% agarose in a 6-well plate. The two layers were covered with medium and incubated at 37°C in 5% CO_2_ for 2 to 4 weeks or until visible colonies appeared. Colonies were stained with 0.01% crystal violet in 10% ethanol and counted in 10 random fields per well.

Cell death was measured after plating 2 × 10^4^ cells in the wells of 96-well plates in triplicate. The next day, the medium was removed and 20 μL lysis buffer (50 mM HEPES, 0.1% CHAPS, 2 mM dithiothreitol, 0.1% Nonidet P-40, 1 mM EDTA, and 1% protease inhibitor) and 20 μL caspase assay buffer (100 mM HEPES, 10% Sucrose, 0.1% CHAPS, 1 mM EDTA, 2 mM dithiothreitol, and 50 μM Caspase-3 substrate) were added. Fluorescence (excitation 380 nm, emission 430 – 460 nm) was measured at 0, 5, and 24 h. DNA replication was measured 24 h after plating of 5×10^4^ cells per well of a 6 well plate by incubation of cells with 15 μM EdU for 2 h. General transcription was measured 24 h after plating of cells by incubation with 1 mM of EU for 40 min or 3 h at 37°C.

All flow cytometry was measured on the BD LSRII machine using FACSDiva software and analyzed using WinList software.

The MNase assay was performed as described (Safina et al. 2017) 24 h after the completion of the 4-OHT treatment (2 × 10^7^ cells per condition). For each reaction, 10 μl DNA was submitted for Bioanalyzer QC analysis.

For the FRAP assay, cells were plated into 35-mm glass bottom plates (Mattek Corp., cat# P35G-1.0-14-C). The assay was performed 24 h after plating using a Leica DMi8 inverted microscope and TCS SP8 laser scanner with Leica Application Suite X (LAS-X) acquisition software (four pre-bleach images with 433 ms intervals and eight bleaches with 100% 522 nm laser power followed by 250 post-bleach measurements every second). Data were accumulated for 15 to 20 cells. All measurements were normalized by the average pre-bleach fluorescence intensity.

#### Immunoblotting

Standard immunoblotting methods were used. Antibodies and dilutions are listed in Table S1.

#### Immunofluorescence staining

Cells were fixed with 4% paraformaldehyde, and staining was performed as previously described (Ref). The antibodies are listed in Table 1. Images were acquired using a Zeiss Axio Observer A1 inverted microscope with an N-Achroplan 100×/1.25 oil lens and a Zeiss MRC5 camera with AxioVision Rel.4.8 software. Image analysis and quantitation were done using ImageJ.

#### RT-qPCR

Total RNA was isolated using TRIzol reagent. First-strand cDNA (30 μL) was synthesized from 500 ng RNA using the SuperScript™ First-Strand Synthesis System. qPCR was performed using 1 μl first-strand cDNA with primers and master mix purchased from Applied Biosystems and the default parameters of the 7900HT sequence detection system (ABI PRISM; Applied Biosystems). To compare gene expression levels between samples, the threshold cycle (CT) value was normalized using the mean CT for the reference gene, GAPDH. The normalized mRNA levels were defined as ΔCT = CT (mean for test gene) – CT (mean for the reference gene). The final data were expressed as the fold difference between the test sample and the control sample, which was defined as 2^-(ΔCT treated with 4-OHT – ΔCT control)^.

### High Throughput Whole Genome Methods

#### RNA-Sequencing

For the RNA sequencing data shown in Figures 3, 4, 8A, S7-S9, and S11 (cells designated #3), and S19, RNA was isolated using Monarch Total RNA Kit (T2010S, New England BioLabs). The total RNA quality was assessed using an Agilent Bioanalyzer. RNA with an overall RIN score >9 was used. RNA was depleted of ribosomal transcripts using the RiboErase kit (Roche). RNA libraries were prepared from 500 ng RNA using the Kapa RNA HyperPrep kit (Roche). All RNA libraries were sequenced using massively parallel sequencing (Illumina, NovaSeq) with 100 base pair paired-end reads. Two independent RNA-seq experiments were performed.

For the RNA sequencing data shown in Figures 8 B and C, S10, and S11 (cells designated #2), RNA was isolated using TRIZOL reagent (Invitrogen, Life Technologies). The total RNA quality was assessed using an Agilent Bioanalyzer. RNA with an overall RIN score >9 was used. RNA was depleted of ribosomal transcripts using the RiboZero Gold kit (Illumina). RNA libraries were prepared from 1000 ng RNA using TruSeq Stranded RNA kit (Illumina). All RNA libraries were sequenced using massively parallel sequencing (Illumina, NextSeq) with 75 base pair single-end reads. Two independent RNA-seq experiments were performed.

#### RNA-Sequencing of human HT1080 cells

HT1080 cells were infected with lentiviruses produced using psi-LVRU6MP encoding shRNA to SSRP1, clone HSH017741-8-LVRU6MP(OS396821) (cat no. CS-HSH0177741 −8-LVRU6MP) or control clone CSHCTR001-LVRU6MP(OSNEG20) (cat no. CSHCTR001-LVRU6MP) from GeneCopoeia. Cells were selected with puromycin for three days and then counted. RNA was isolated using the Monarch Total RNA Kit (T2010S, New England BioLabs). A 1:1000 dilution of ERCC RNA Spike-in Mix1 (Life Technologies) was added to 100 ng total RNA at a ratio corresponding to the number of input cells used for RNA extraction. RNA was depleted of ribosomal transcripts using the RiboZero Gold kit (Illumina). RNA libraries were prepared from 1 μg total RNA using TruSeq Stranded Total RNA kit (Illumina) according to the manufacturer’s instructions. The resulting pool was loaded into the appropriate NextSeq Reagent cartridge, for 75 single-end sequencing, and sequenced using the NextSeq500 following the manufacturer’s recommended protocol (Illumina).

#### Chromatin Immunoprecipitation (ChIP) and sequencing

ChIP-Seq samples were prepared from mouse cells using the SimpleChIP Kit (cat no. 9003, Cell Signaling Technology). Immunoprecipitation was performed using the mouse monoclonal 10D1 anti-SSRP1 antibody (10 μg/IP; cat no. 609702, BioLegend, Inc). The histone H3(D2B12)XP rabbit monoclonal antibody (4620) provided in the kit was used as a positive control. DNA isolated after MNase digestion was used as the input DNA.

For the ChiP-Seq, 2 ng chromatin-immunoprecipitated DNA was used to generate the library for next-generation sequencing using the ThruPLEX DNA seq kit (Rubicon Genomics, Inc.) according to the manufacturer’s instructions. The DNA libraries were quantitated using the KAPA Biosystems qPCR kit and pooled in an equimolar fashion. Each pool was denatured and diluted to 2.4 pM with 1% PhiX control library. The resulting pool was loaded into the appropriate NextSeq Reagent cartridge for 75 paired-end sequencing and sequenced on a NextSeq500 following the manufacturer’s recommended protocol (Illumina).

#### Analyses of NGS data

Raw reads that passed the quality filter from Illumina Real Time Analysis (RTA) software were mapped to the latest reference genome (hg38 for human and mm10 for mouse samples, respectively) using Tophat2 (Trapnell et al. 2009). The gene expression quantitation was generated using the Subsread package (Liao et al. 2014) with GenCode for differential expression analysis using DESeq2 (Love et al. 2014). Pathway analysis was done using GSEA (Subramanian et al. 2005) with C2 curated gene sets in MSigDB. ChIP-Seq reads were mapped to reference genomes using bwa (Li and Durbin 2009), and the narrow peaks were identified by MACS2 (Zhang et al. 2008) using the input DNA as a control. Because SSRP1 is a protein that occupies the entire gene, we only used non-overlapping protein-coding genes (with NM prefix in RefGene) to study the relationship of gene expression with SSRP1 coverage. Genes were grouped into different categories according to the RPKM/FPKM values generated using the edgeR (Robinson et al. 2010) Bioconductor R package. The big wiggle files and SSRP1 profiles for the whole gene and around the TSS were generated using the deepTools suite (Ramirez et al. 2016). The correlation coefficients and p-values between RNA expression and SSRP1 coverage were calculated using R statistical software. For human RNA-Seq samples with ERCC spike-in, the normalization factors were determined using the loess normalization function from affy (Gautier et al. 2004) Bioconductor package before differential gene analysis using DESeq2.

#### Animal experiments

All animal experiments were conducted according to a protocol approved by the Institute Animal Care and Use Committee at Roswell Park Comprehensive Cancer Center. The facility has been certified by the American Association for Accreditation of Laboratory Animal Care in accordance with the current regulations and standards of the U.S. Department of Agriculture and the U.S. Department of Health and Human Services.

Generation of *Ssrp1^fl/fl^; CreER^T2^* mice and isolation of mouse skin fibroblasts from tails of these mice were described (Sandlesh et al. 2018).

For *in vivo* tumor growth, 1×10^6^ cells (Im or Tr #3) were injected subcutaneously into both lateral flanks of ten 6-week-old female SCID (C.B-Igh-1^b^/IcrTac-Prkdc^scid^) mice (n = 20). In mice inoculated with immortalized cells, only one tumor appeared during three months of observation. Tumors were visible in most Tr-inoculated mice 3 to 5 days after inoculation. Mice were randomly divided into treatment and control groups (n = 10) 48 h post-inoculation. The treatment and control groups received 1 mg/100μL tamoxifen or 5% ethanol i.p. following a 3 days on/1 day off schedule, respectively. Treatment was continued until the control tumors reached 1 cm^3^.

#### Statistical analysis

Data were compared between the control and treated groups using the unpaired t-test (Mann-Whitney test). Analyses were conducted using GraphPad Prism 7.03 and all p-values were two-sided.

## Supporting information

## Acknowledgments

We are grateful to Prashant Singh from Genomics Shared Resource of Roswell Park Comprehensive Cancer for always available help with processing of our sequenceing needs, to Carlos Cedena from Flow and Imaging Shared Resource of Roswell Park Comprehensive Cancer for the help with confocal microscopy and FRAP, to Caterine Burkhart from Burkhart Documents Solutions for critical reading and editing of the manuscript. Funding: National Cancer Institute: R01CA197967: Dr. Katerina V. Gurova; National Cancer Institute: R21CA198395: Dr. Katerina V. Gurova; National Cancer Institute: P30CA016056: to Roswell Park Comprehensive Cancer Center; Susan G. Komen (US): CCR13264604: Dr. Katerina V, Gurova; Incuron LLC: Dr. Katerina V. Gurova.

## Author Contributions

Poorva Sandlesh – design and execution of most of experiments, writing and editing of the manuscript, Alfiya Safina – preparation of samples for ChIP-sequencing, Imon Goswami – isolation of cells from animals, MNAse digestion experiments, Laura Prendergust – analysis of DNA damage in cells, Spenser Rosario – analysis of RNA-seq data, Eduardo C Gomez – analysis of RNA-seq data, Jianmin Wang – all bioinformatics processing and analyses, Katerina V Gurova – development of the concept of the study, design and analyses of data, writing and editing of the manuscript.

