## Supplementary material for "Prevention of chromatin destabilization by FACT is crucial for malignant transformation"

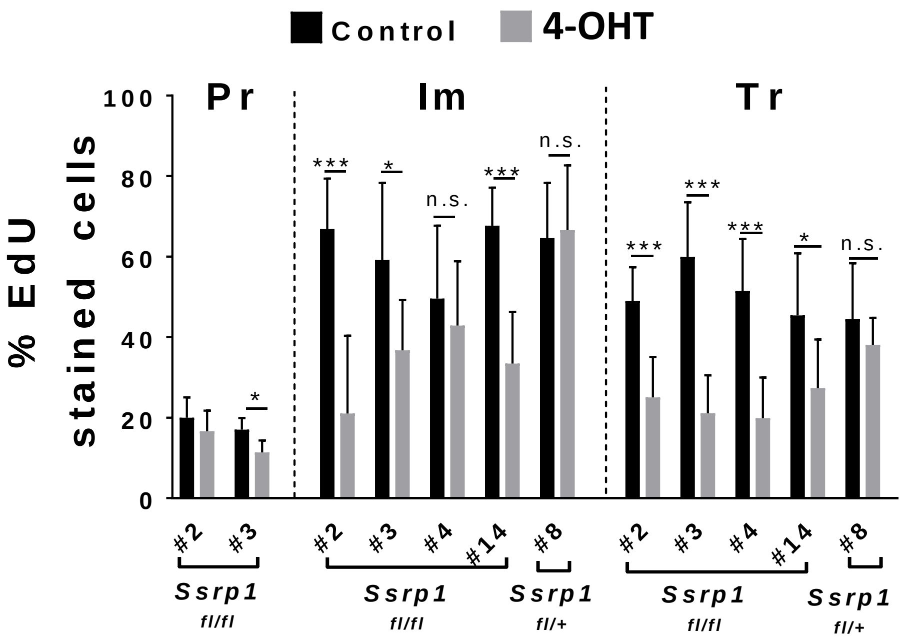


**Figure S4. Effect of *Ssrp1* KO on EdU incorporation in Pr, Im, and Tr cells.** The cell proliferation assay was performed using *Ssrp1*$^{fl/fl}$ *CreER*$^{T2+/+}$ (clones: #2, #3, #4, #14) and *Ssrp1*$^{fl/+}$ *CreER*$^{T2+/+}$ (clone #8) fibroblasts. The effects of *Ssrp1* KO (grey bar) are presented relative to the control (black bar) for each cell line. * <0.01, **<0.001 and ***<0.0001
