## Supplementary material for "Prevention of chromatin destabilization by FACT is crucial for malignant transformation"

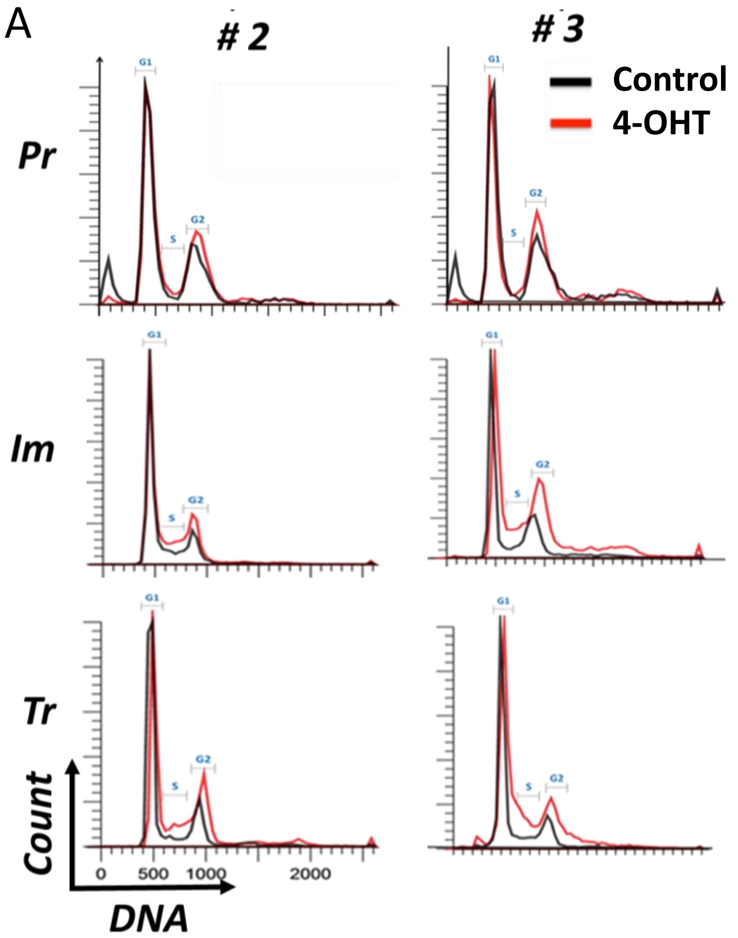

B

| Genotype | Phenotype | Cell # | G0 Control | G0 + 4OHT (2μM) | G1 Control | G1 + 4OHT (2μM) | S Control | S + 4OHT (2μM) | G2/M Control | G2/M + 4OHT (2μM) |
|---|---|---|---|---|---|---|---|---|---|---|
| *Ssrp1* $^{fl/fl}$ *CreER* $^{T2+/+}$ | Primary | #2 | 5.57 | 0.82 | 51.28 | 44.39 | 2.35 | 3.32 | 27.92 | 32.27 |
| | | #3 | 7.92 | 1.08 | 45.92 | 41.85 | 2.37 | 5.36 | 26.16 | 33.54 |
| | Immortalized | #2 | 0.01 | 0.05 | 62.61 | 51.34 | 10.33 | 15.57 | 22.64 | 37.85 |
| | | #3 | 0.07 | 0.33 | 54.04 | 29.02 | 7.64 | 15.64 | 25.45 | 37.12 |
| | Transformed | #2 | 0.03 | 0.14 | 68.23 | 45.04 | 10.17 | 22.52 | 17.79 | 29.12 |
| | | #3 | 0.19 | 0.66 | 59.68 | 42.13 | 12.78 | 24.91 | 20.78 | 35.13 |
| *Ssrp1* $^{fl/+}$ *CreER* $^{T2+/+}$ | Immortalized | #8 | 0.07 | 0.23 | 44.27 | 46.86 | 21.92 | 20.02 | 28.56 | 28.8 |
| | Transformed | #8 | 0.11 | 0.14 | 59.44 | 56.33 | 12.98 | 13.34 | 13.4 | 15.09 |

C

| | | Aver Control | Aver + 4OHT (2μM) | T test | SDV Control | SDV + 4OHT (2μM) |
|---|---|---|---|---|---|---|
| G1 | Primary | 48.6 | 43.12 | 0.080162 | 2.68 | 1.27 |
| | Immortalized | 58.325 | 40.18 | 0.115285 | 4.285 | 11.16 |
| | Transformed | 63.955 | 43.585 | 0.043788 | 4.275 | 1.455 |
| S | Primary | 2.36 | 4.34 | 0.150145 | 0.01 | 1.02 |
| | Immortalized | 8.985 | 15.605 | 0.065418 | 1.345 | 0.035 |
| | Transformed | 11.475 | 23.715 | 0.002861 | 1.305 | 1.195 |
| G2/M | Primary | 27.04 | 32.905 | 0.080464 | 0.88 | 0.635 |
| | Immortalized | 24.045 | 37.485 | 0.04168 | 1.405 | 0.365 |
| | Transformed | 19.285 | 32.125 | 0.037262 | 1.495 | 3.005 |

**Figure S3. Effect of *Ssrp1* KO on the cell cycle distribution in primary (Pr), immortalized (Im), and transformed (Tr) cells.** A. Cell cycle distribution profiles for *Ssrp1* $^{fl/fl}$ *CreER* $^{T2+/+}$ cells treated with 4-OHT (red) or vehicle (black). B. Quantitation of cell cycle distribution data using ModFit. C. Statistical evaluation of the cell cycle distribution data.
