## Supplementary material for "Prevention of chromatin destabilization by FACT is crucial for malignant transformation"

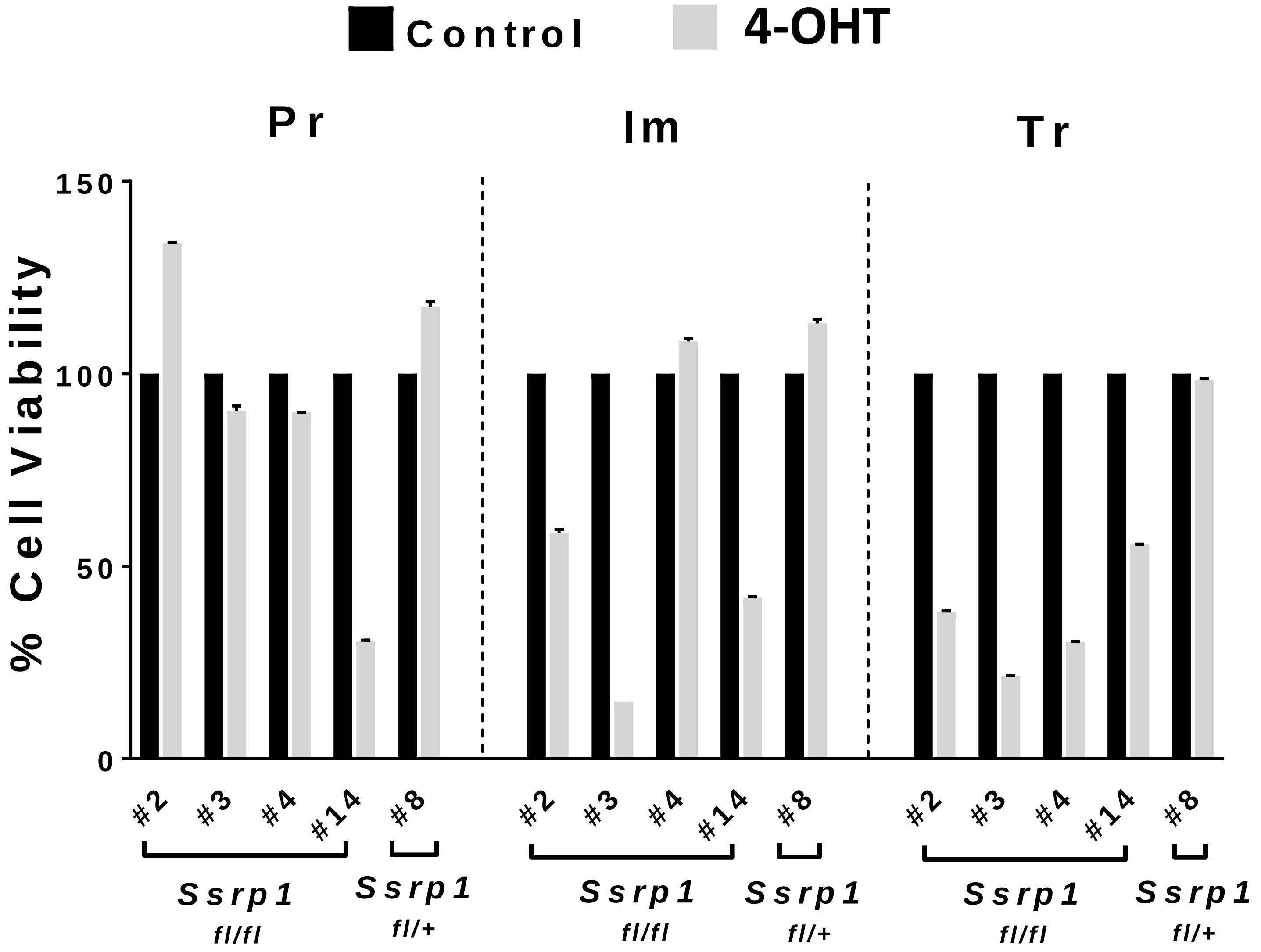


**Figure S2. Viability of primary (Pr), immortalized (Im), and transformed (Tr) MSFs before and after *Ssrp1* KO.** Cells were plated following a 5-day treatment with 4-OHT or vehicle. Cell viability was assessed using a resazurin-based assay 48 h after plating. Data are presented as the mean ± SD (n = 3).
