## Supplementary material for "Prevention of chromatin destabilization by FACT is crucial for malignant transformation"

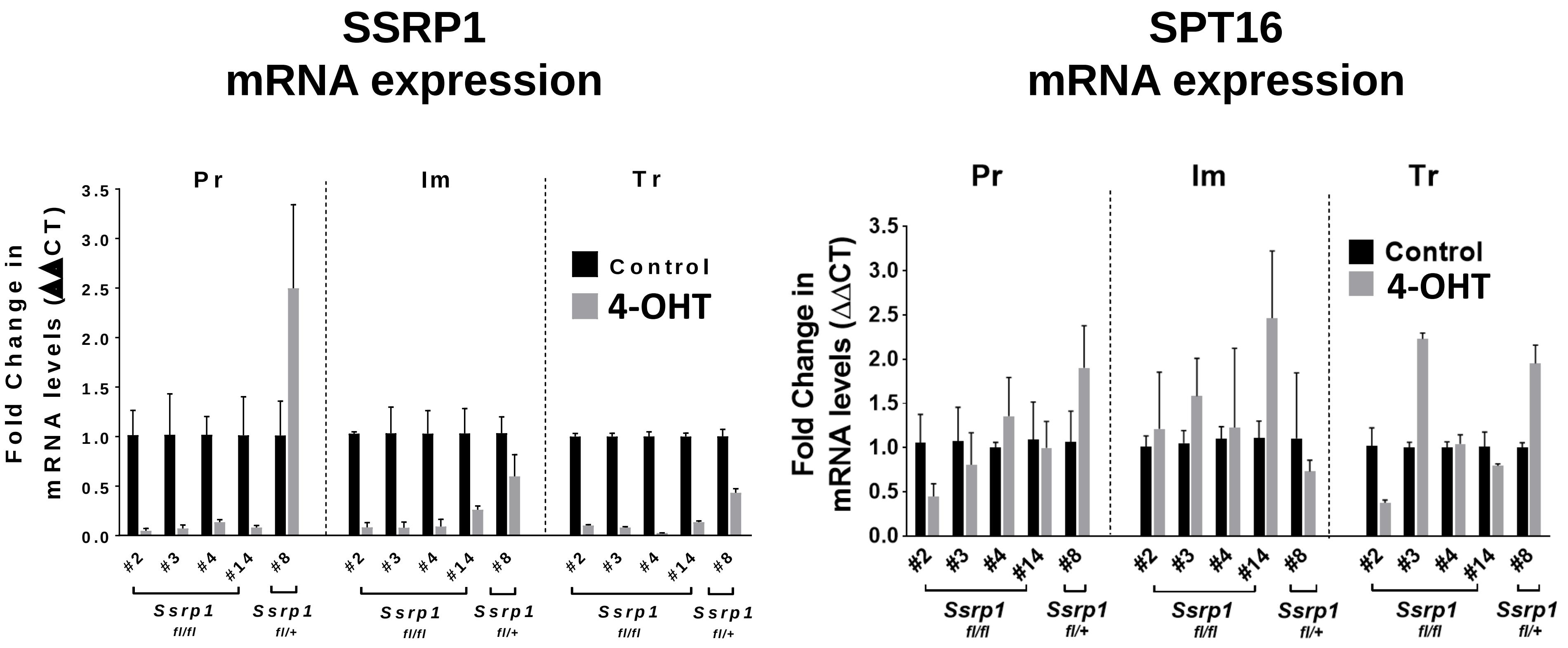


**Figure S1. mRNA levels of *Ssrp1* and *Supt16*** (gene encoding SPT16) in primary (Pr), immortalized (Im) and transformed (Tr) MSFs from *Ssrp1*$^{fl/fl}$ *CreER*$^{T2+/+}$ (#2, #3, #4, #14) and *Ssrp1*$^{fl/+}$ *CreER*$^{T2+/+}$ (#8) mice. The MSFs were treated with 4-OHT (grey bar) for five days. The expression levels were measured using qRT-PCR and are presented relative to the untreated cells (black bar). Data are presented as the mean ± SD (n = 3).
