## Supplementary material for "Prevention of chromatin destabilization by FACT is crucial for malignant transformation"

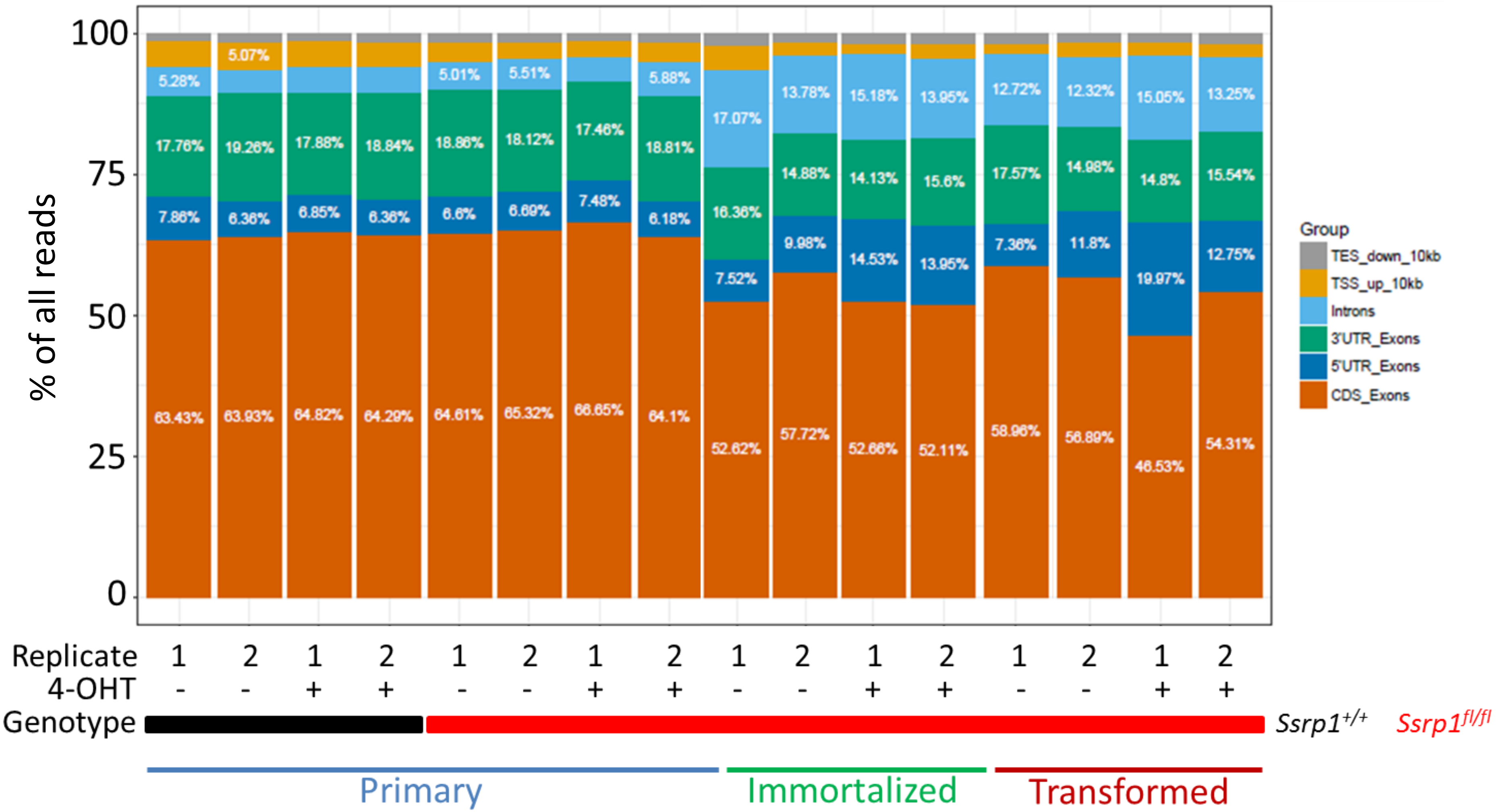


**Figure S19. Distribution of RNA-seq reads among genomic regions associated with genic features.** TES_down_10kb, read located within 10 kB downstream of the transcription end site; TSS_up_10kb, 10 kb upstream of the transcription start site; introns, within the intron of any gene; 3'UTR_Exons, within the last exon of a gene; 5'UTR_Exon, within the first exon of a gene; CDS_Exons, within all other exons of a gene. Total RNA was isolated from replicate cultures of 4-OHT- or vehicle-treated primary (*Ssrp1* wild type #11, floxed # 4), immortalized (#3), and transformed (#3) cells using the NEB Monarch RNA isolation kit. The library was prepared using the TruSeq Stranded Total RNA kit (Illumina Inc) and was used for 100 bp paired-end sequencing in Ia llumina  NovaSeq flow cell.
