## Supplementary material for "Prevention of chromatin destabilization by FACT is crucial for malignant transformation"

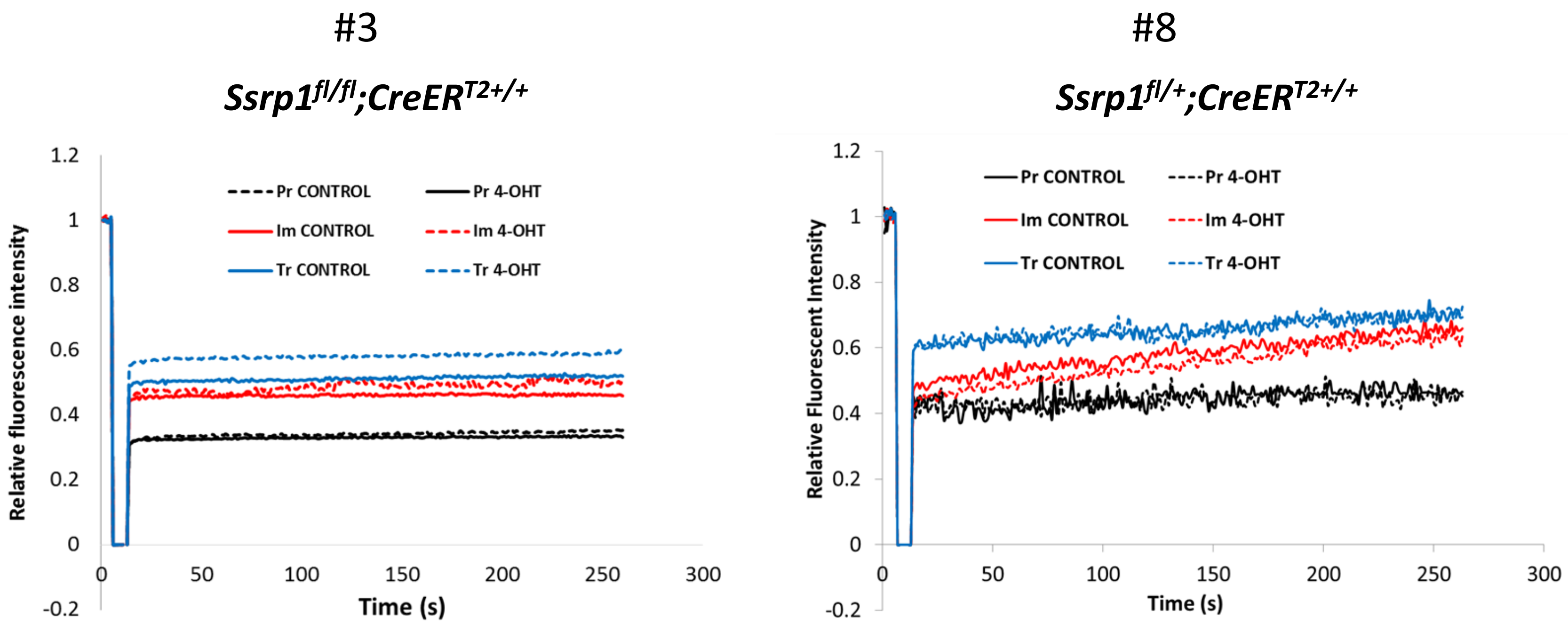


**Figure S18. FRAP assay for primary (Pr), immortalized (Im), and transformed (Tr) cells of two genotypes following treatment with 4-OHT or vehicle for five days.** After treatment, the cells were plated in glass-bottom plates and FRAP was measured in 15-20 cells of each type using the same settings and conditions. Each curve represents the average for all measured cells normalized by the average intensity of fluorescence before bleaching (five measurements at 400 ms intervals). Recovery was measured every second during a 250-second period.
