## Supplementary material for "Prevention of chromatin destabilization by FACT is crucial for malignant transformation"

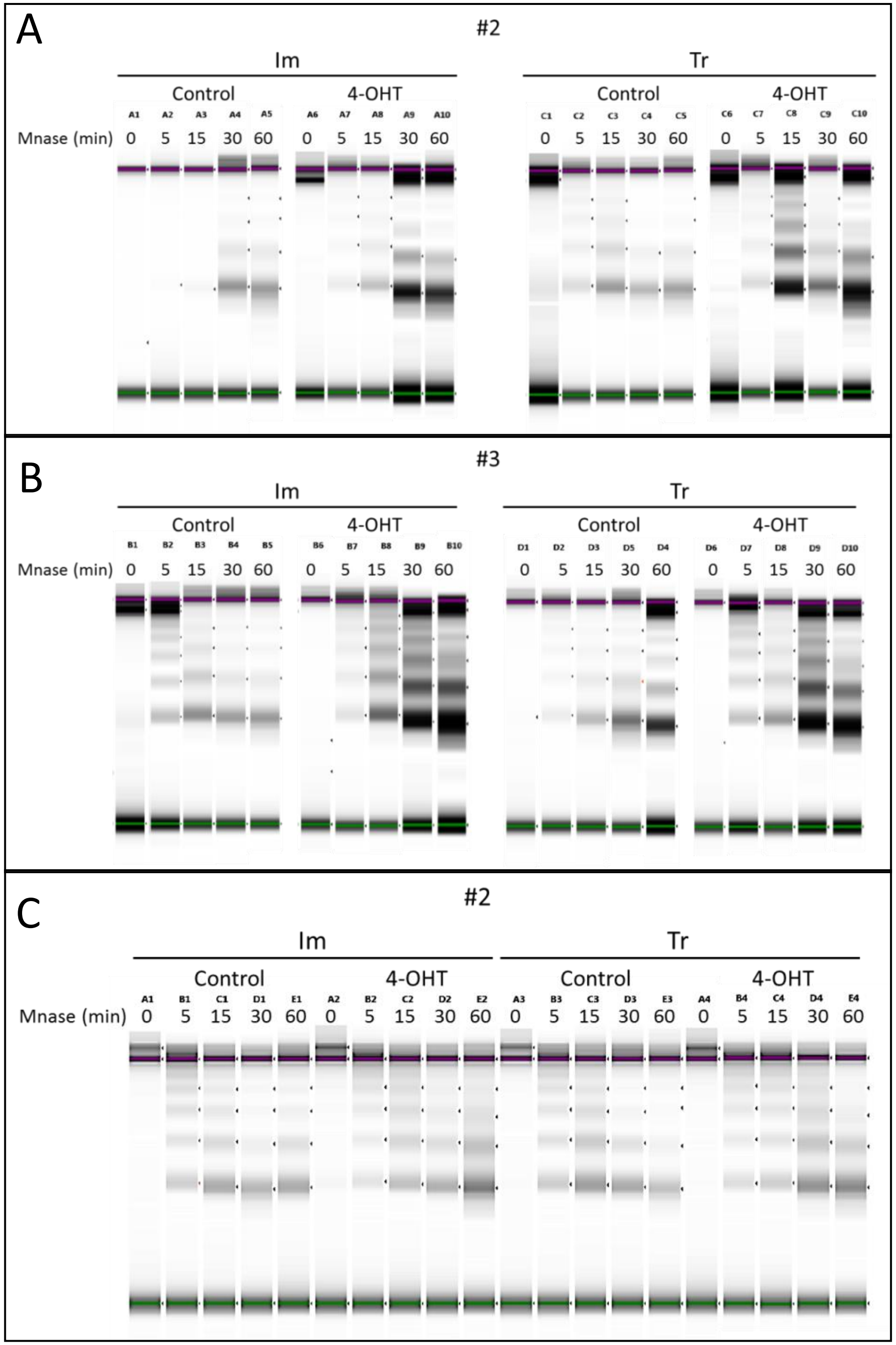


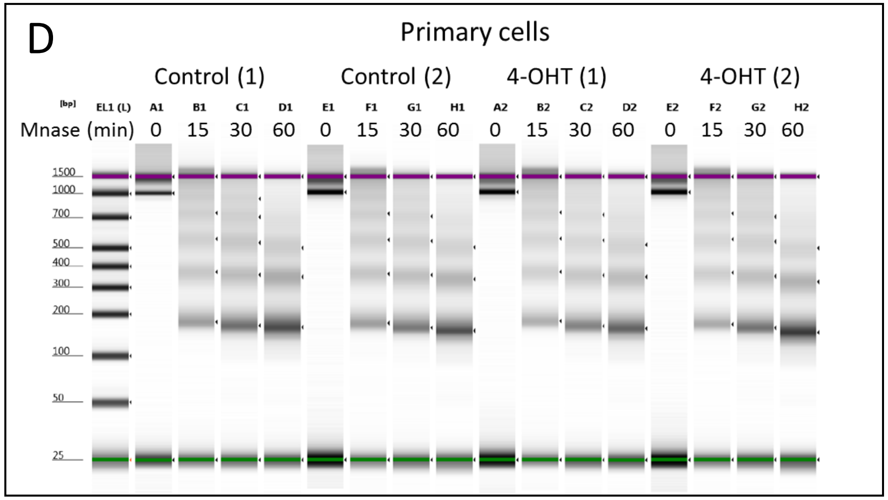


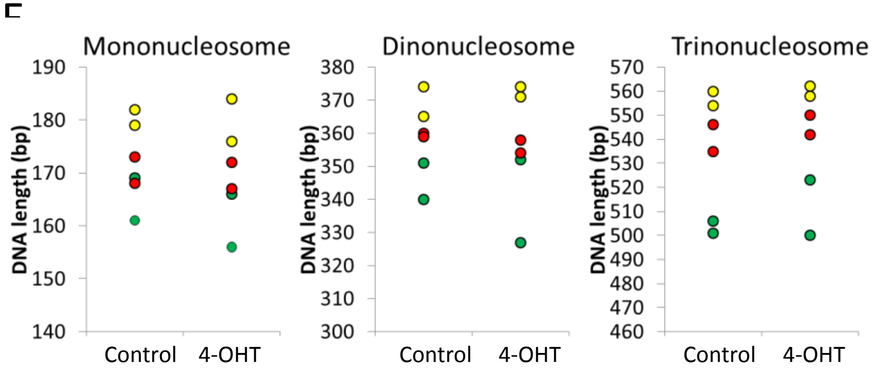


**Figure S17. MNase digestion of nuclei from cells with floxed *Ssrp1* treated with 4-OHT or vehicle for five days.** A, B, C represent three independent experiments with immortalized (Im) and transformed (Tr) cells performed 24 h after the completion of 4-OHT treatment. . The same number of nuclei was used for each condition. D. Experiment done with primary cells (in two replicates). Isolated DNA was run on capillary electrophoresis (BioAnalyzer). Purple and green lines indicate positions of spike-in controls. E. Comparison of length of peaks corresponding to mono-, di- and trinucleosomes in primary cells before and after treatment with 4-OHT. Colors correspond to the time of treatment with MNase: yellow - 15 min, red – 30 min, green – 60 min.
