## Supplementary material for "Prevention of chromatin destabilization by FACT is crucial for malignant transformation"

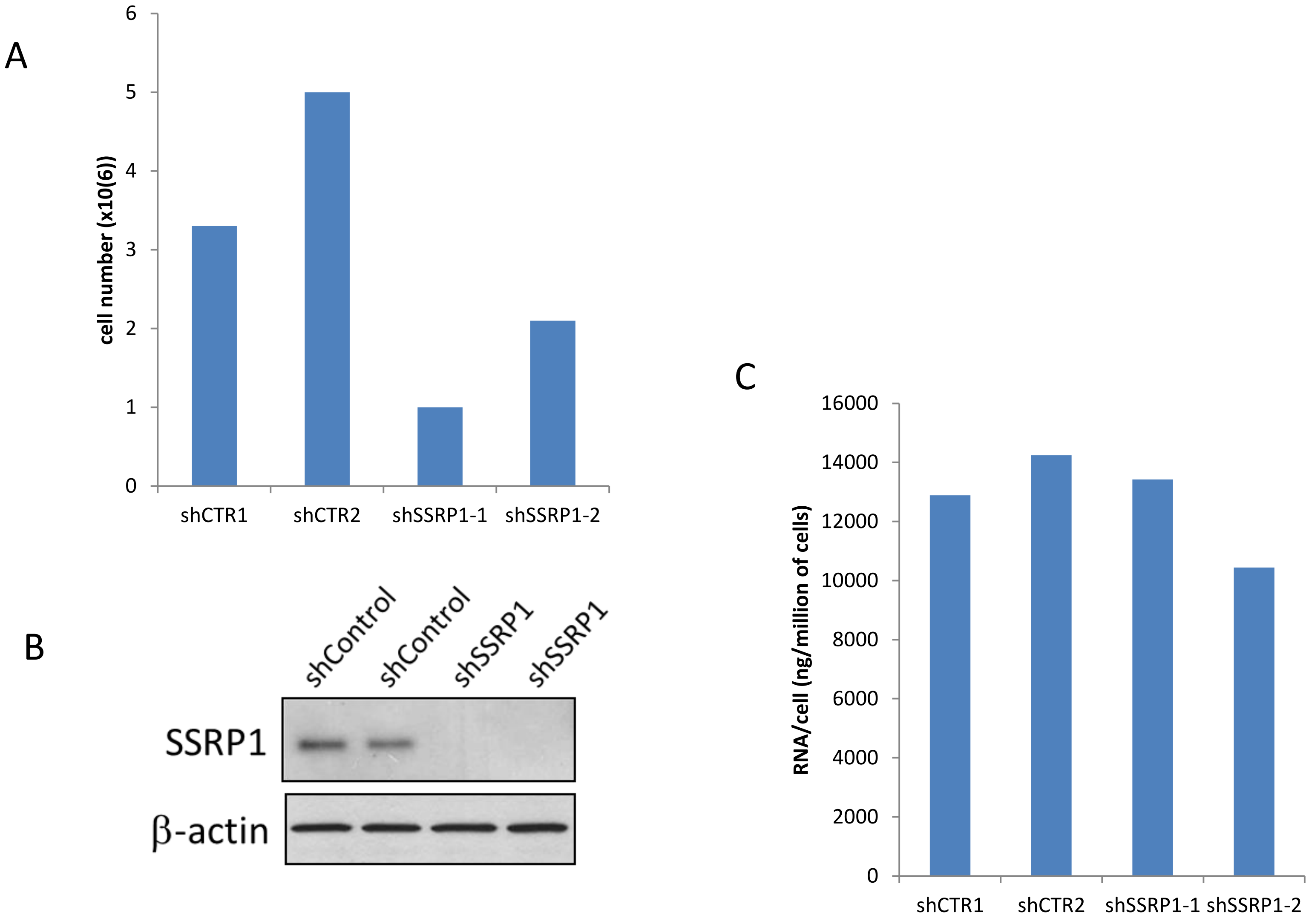


**Figure S16. Characterization of the HT1080 samples used for RNA-seq**. Two plates (1 and 2) with equal numbers of HT1080 cells were transduced with shControl (shCTR) or shSSRP1 lentiviruses. After 72 h, cells were counted (A) and then used for western blotting (B) or RNA isolation (C).
