## Supplementary material for "Prevention of chromatin destabilization by FACT is crucial for malignant transformation"

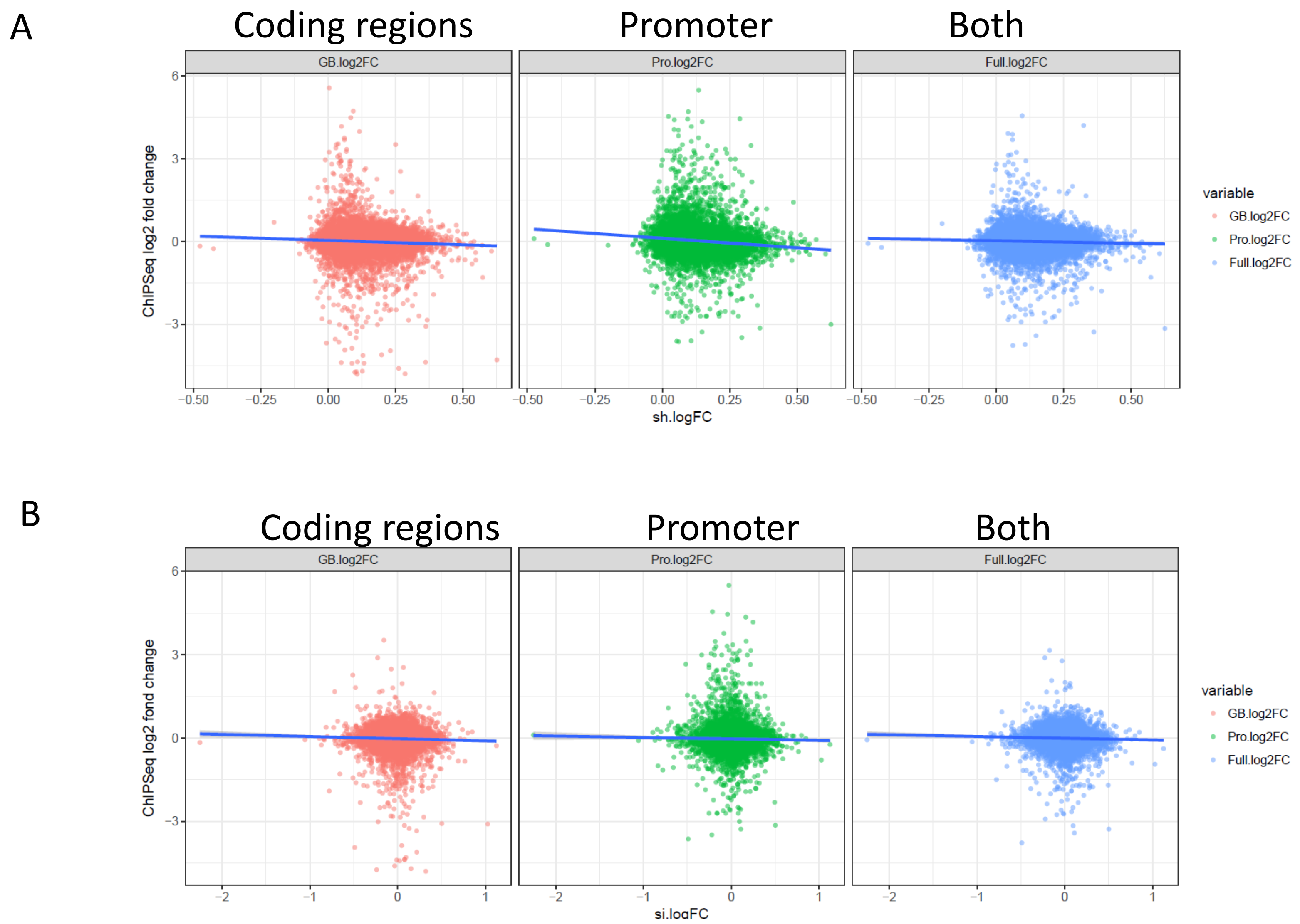


**Figure S15. Correlation between FACT coverage and changes in gene expression following FACT knockdown.** A. HT1080 cells were transduced with lentiviral shRNA to SSRP1 or GFP. Gene expression was measured five days after transduction and selection with puromycin using the Illumina Beads Array. B. HT1080 cells were co-transfected with siRNAs to SSRP1 and SPT16 or control siRNA. Gene expression was measured 72 h after transfection using the Illumina Beads Array. Two independent experiments were performed. In both experiments, analysis was performed separately for the coding regions and promoters as well as both together.
