## Supplementary material for "Prevention of chromatin destabilization by FACT is crucial for malignant transformation"

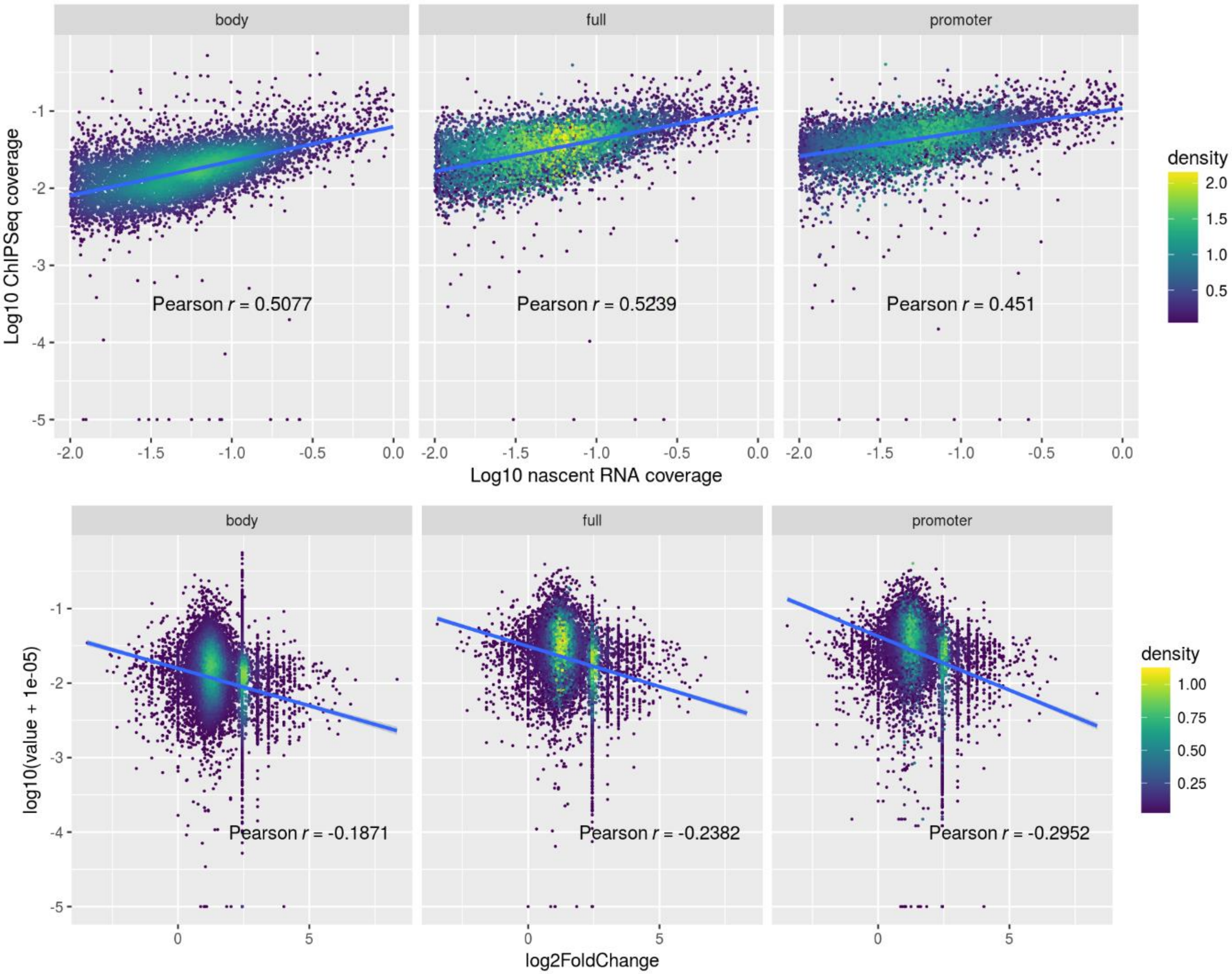


**Figure S14. Correlation between gene transcription and SSRP1 enrichment** in human fibrosarcoma cells (HT1080) analyzed at the coding regions (body), promoter regions (promoter), or both regions (full). A. Correlation between SSRP1 coverage and transcription in basal conditions. B. Correlation between SSRP1 coverage and changes in gene expression in cells transduced with shSSRP1 versus shControl.
