## Supplementary material for "Prevention of chromatin destabilization by FACT is crucial for malignant transformation"

### A Nascent RNA Seq data

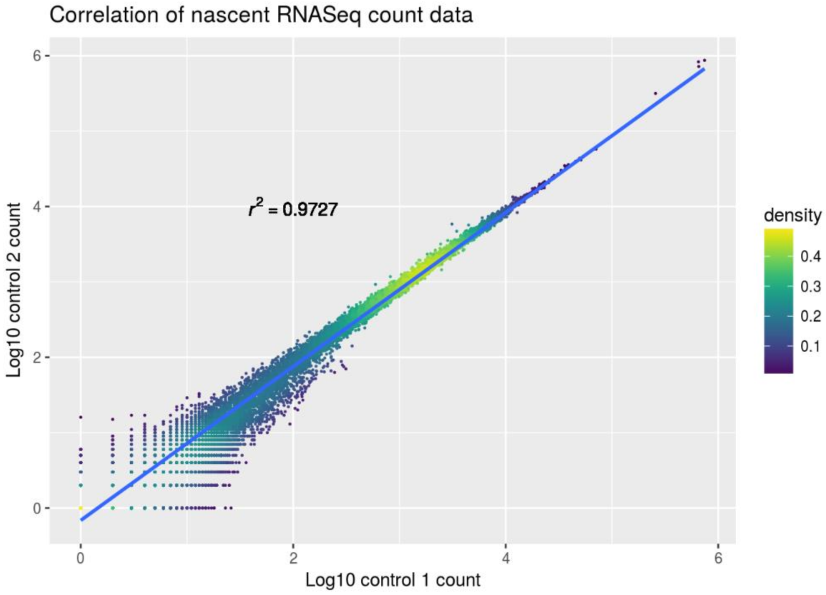


Figure S13. Correlation between replicates of nascent RNA-seq (A) and total RNA-seq (B) data from HT1080 cells transduced with control or SSRP1 shRNA.

### B RNASeq data

For RNASeq data, we have both wild type and SSRP1 KO samples, we will check wild type only.

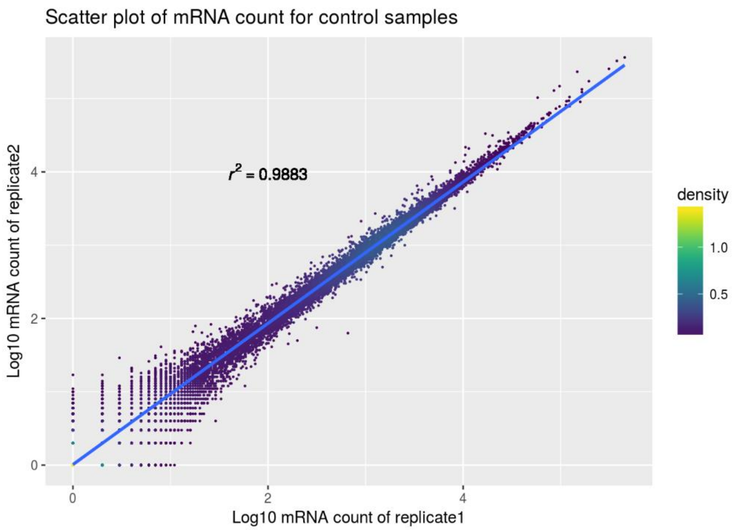


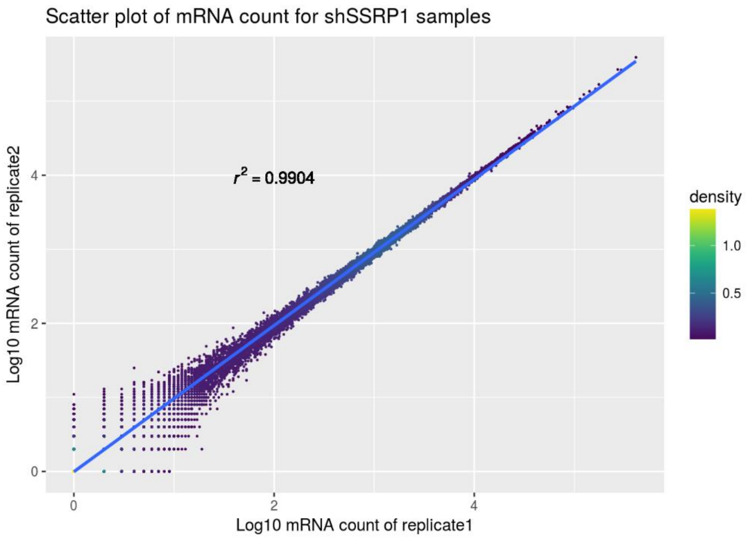
