## Supplementary material for "Prevention of chromatin destabilization by FACT is crucial for malignant transformation"

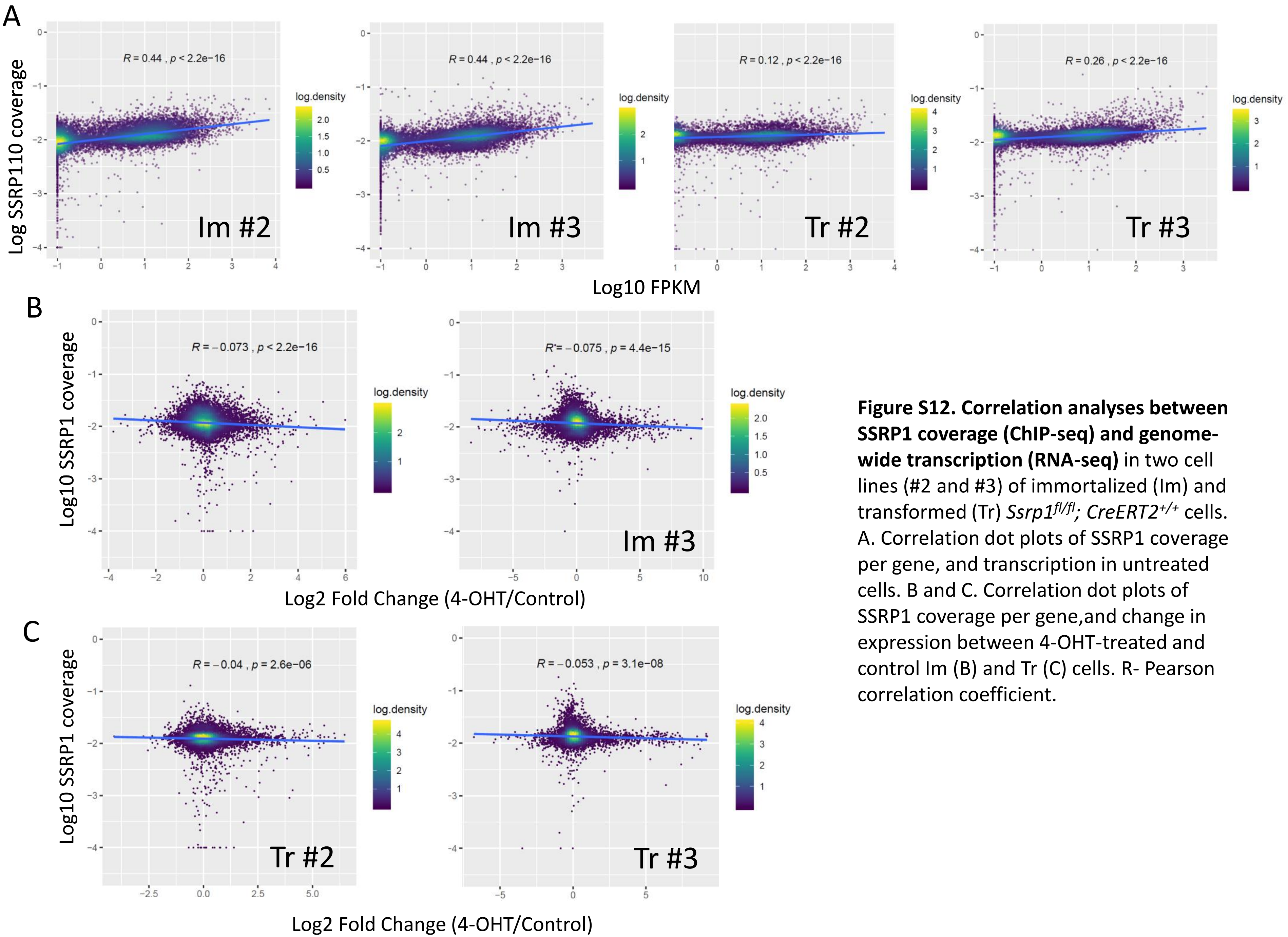


**Figure S12. Correlation analyses between SSRP1 coverage (ChIP-seq) and genome-wide transcription (RNA-seq)** in two cell lines (#2 and #3) of immortalized (Im) and transformed (Tr) *Ssrp1*$^{fl/fl}$*; CreERT2*$^{+/+}$ cells. A. Correlation dot plots of SSRP1 coverage per gene, and transcription in untreated cells. B and C. Correlation dot plots of SSRP1 coverage per gene,and change in expression between 4-OHT-treated and control Im (B) and Tr (C) cells. R- Pearson correlation coefficient.
