## Supplementary material for "Prevention of chromatin destabilization by FACT is crucial for malignant transformation"

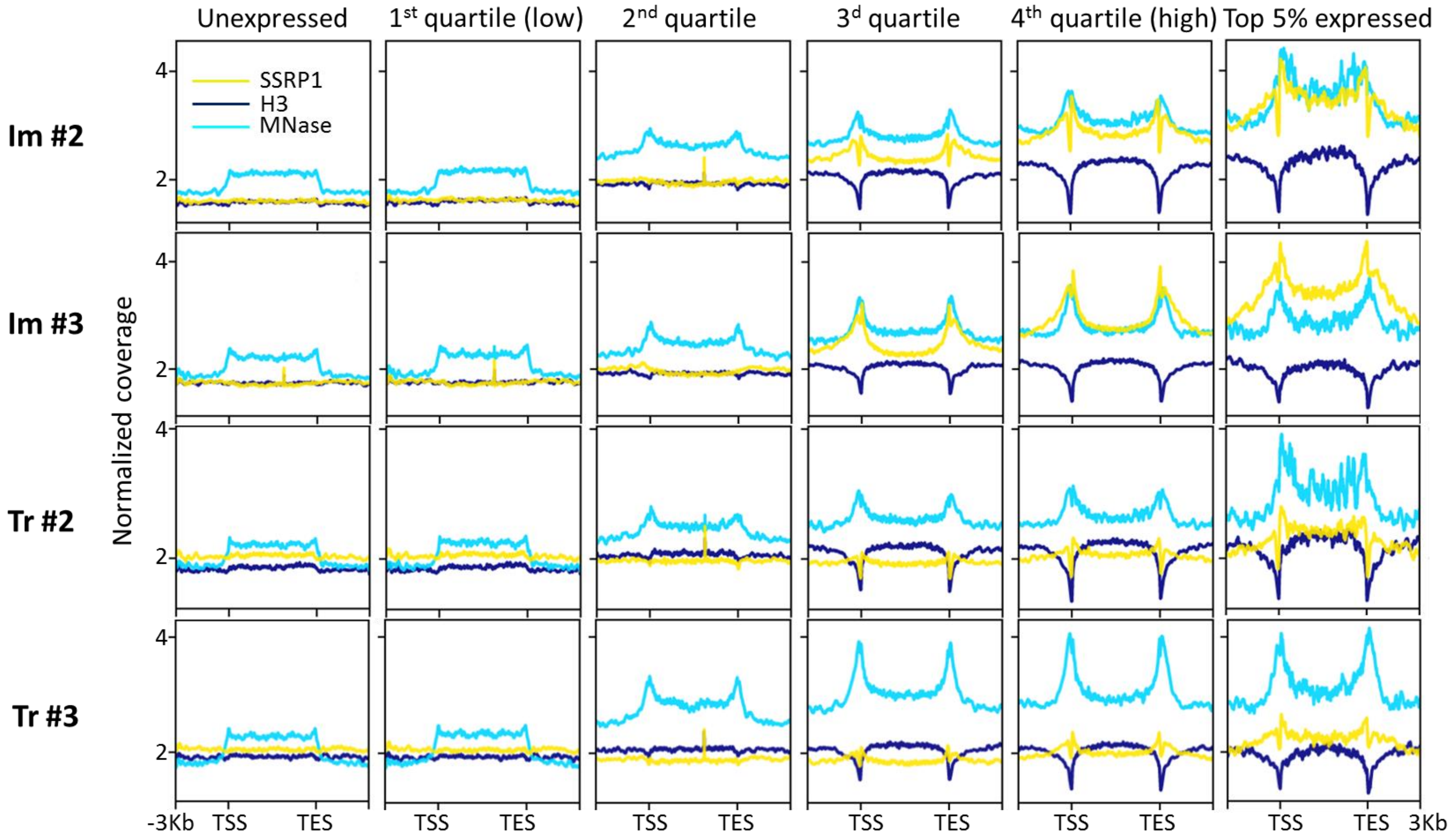


**Figure S11. Average SSRP1 and histone H3 distribution profiles** for two cell lines (#2 and #3) of immortalized (Im) and transformed (Tr) *Ssrp1*$^{fl/fl}$*; CreERT2*$^{+/+}$ MSFs depending on the level of gene expression (single end RNA-seq). The third profile represents the Mnase-digested chromatin used for ChIP.
