## Supplementary material for "Prevention of chromatin destabilization by FACT is crucial for malignant transformation"

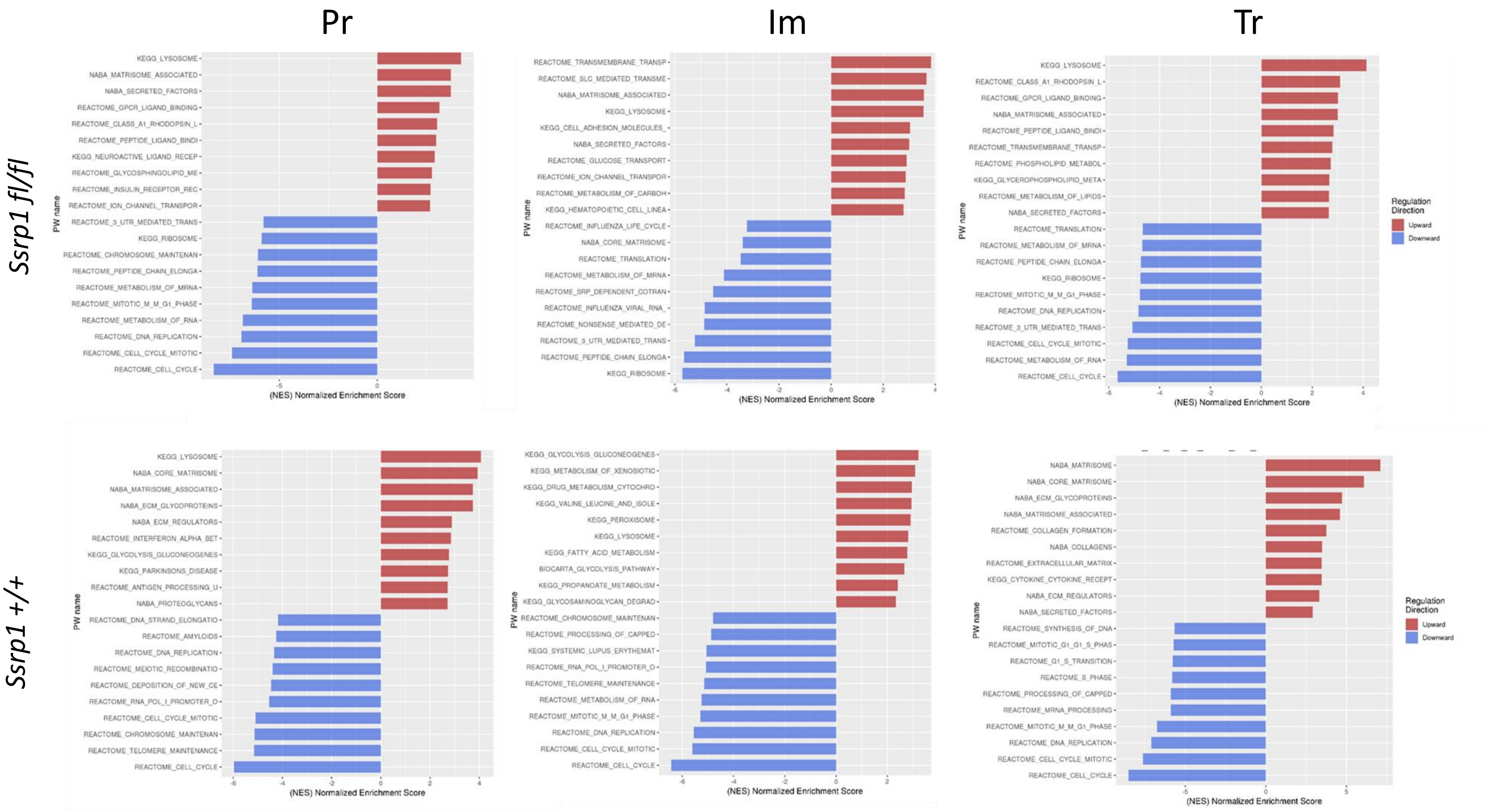


**Figure S10. Gene Set Enrichment Analyses** of genes upregulated (red) or downregulated (blue) in primary (Pr), immortalized (Im), and transformed (Tr) cells from *Ssrp1*$^{+/+}$ *; CreER*$^{T2+/+}$ (bottom row) or *Ssrp1*$^{fl/fl}$ *; CreER*$^{T2+/+}$ (top row) mice in response to 4-OHT treatment.
