## Supplementary material for "Prevention of chromatin destabilization by FACT is crucial for malignant transformation"

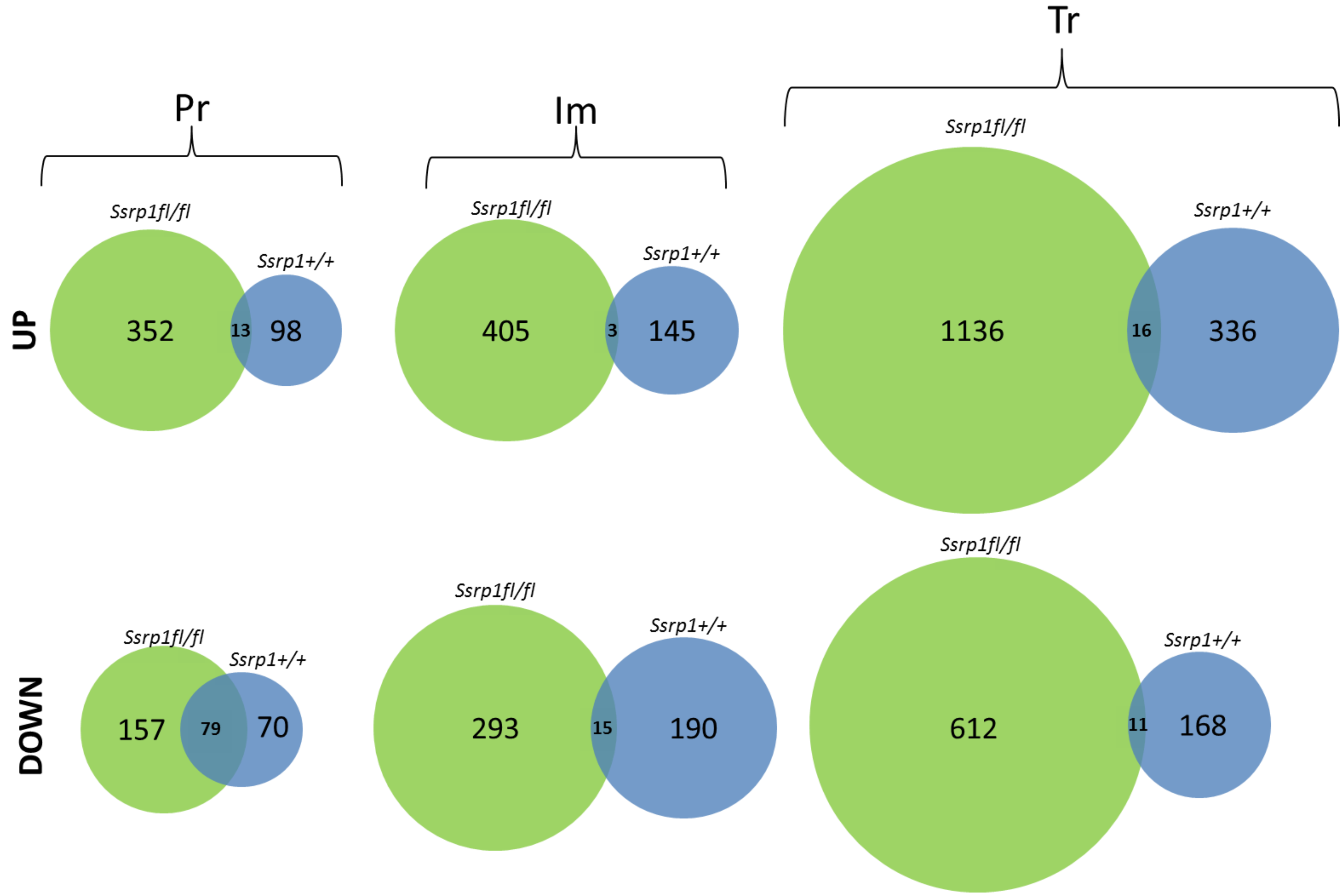


**Figure S9. Venn diagram** showing the number of shared differently up-regulated and downregulated genes (fold change >1.5 and adjusted p-value < 0.05) following 4-OHT treatment in primary (Pr), immortalized (Im), and transformed (Tr) cells from *Ssrp1*$^{+/+}$ *; CreER*$^{T2+/+}$ (blue) and *Ssrp1*$^{fl/fl}$ *; CreER*$^{T2+/+}$ (green) mice.
