## Supplementary material for "Prevention of chromatin destabilization by FACT is crucial for malignant transformation"

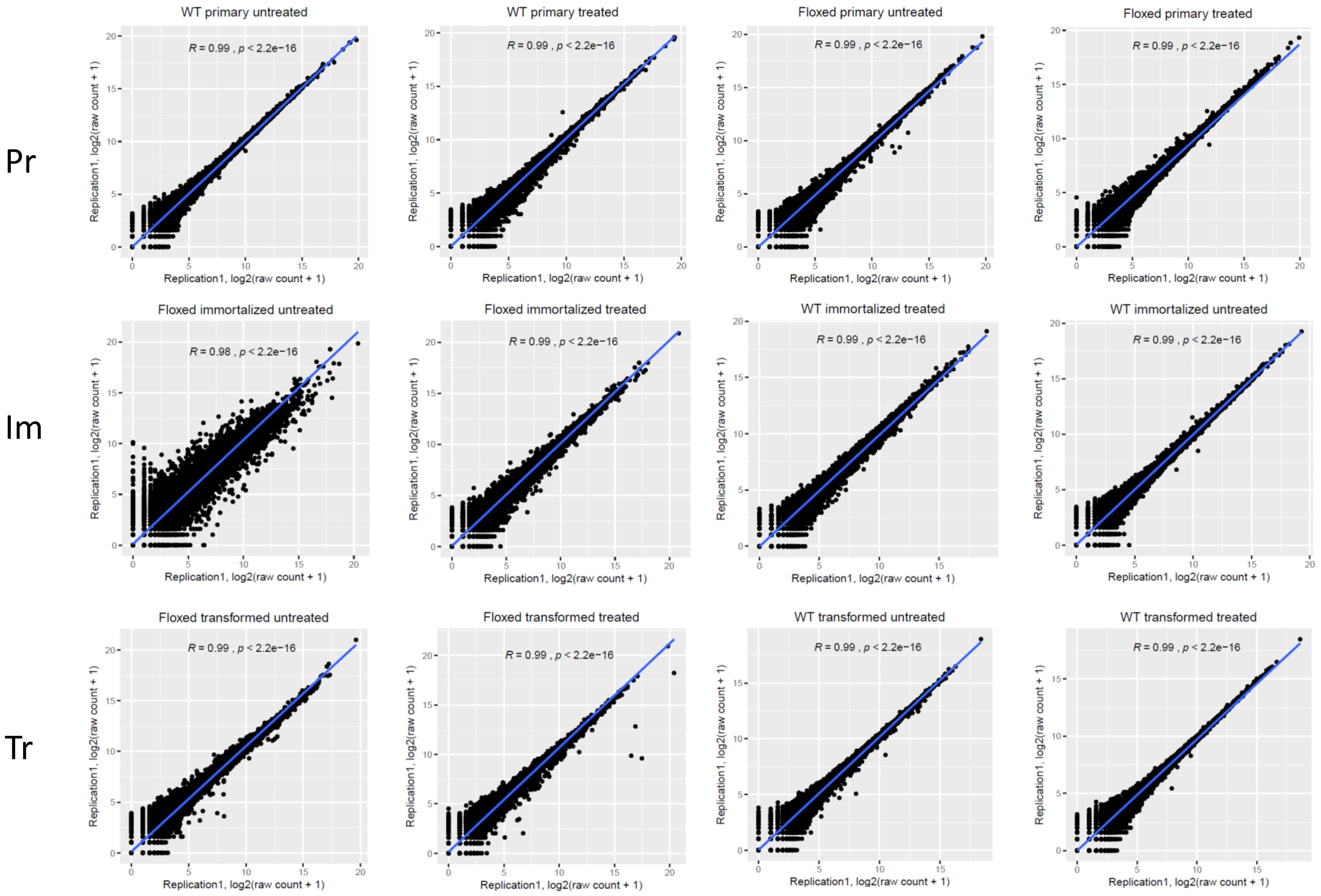


**Figure S8. Correlation dot plots for RNA-seq replicates from mouse cells.** Primary (Pr), immortalized (Im), and transformed (Tr) cells from *Ssrp1*$^{+/+}$ *; CreER*$^{T2+/+}$ (WT) or *Ssrp1*$^{fl/fl}$ *; CreER*$^{T2+/+}$ (floxed) mice were treated with 4-OHT or vehicle for five days. RNA-sequencing was performed for each condition for two replicates. R – Pearson correlation coefficient.
