## Supplementary material for "Prevention of chromatin destabilization by FACT is crucial for malignant transformation"

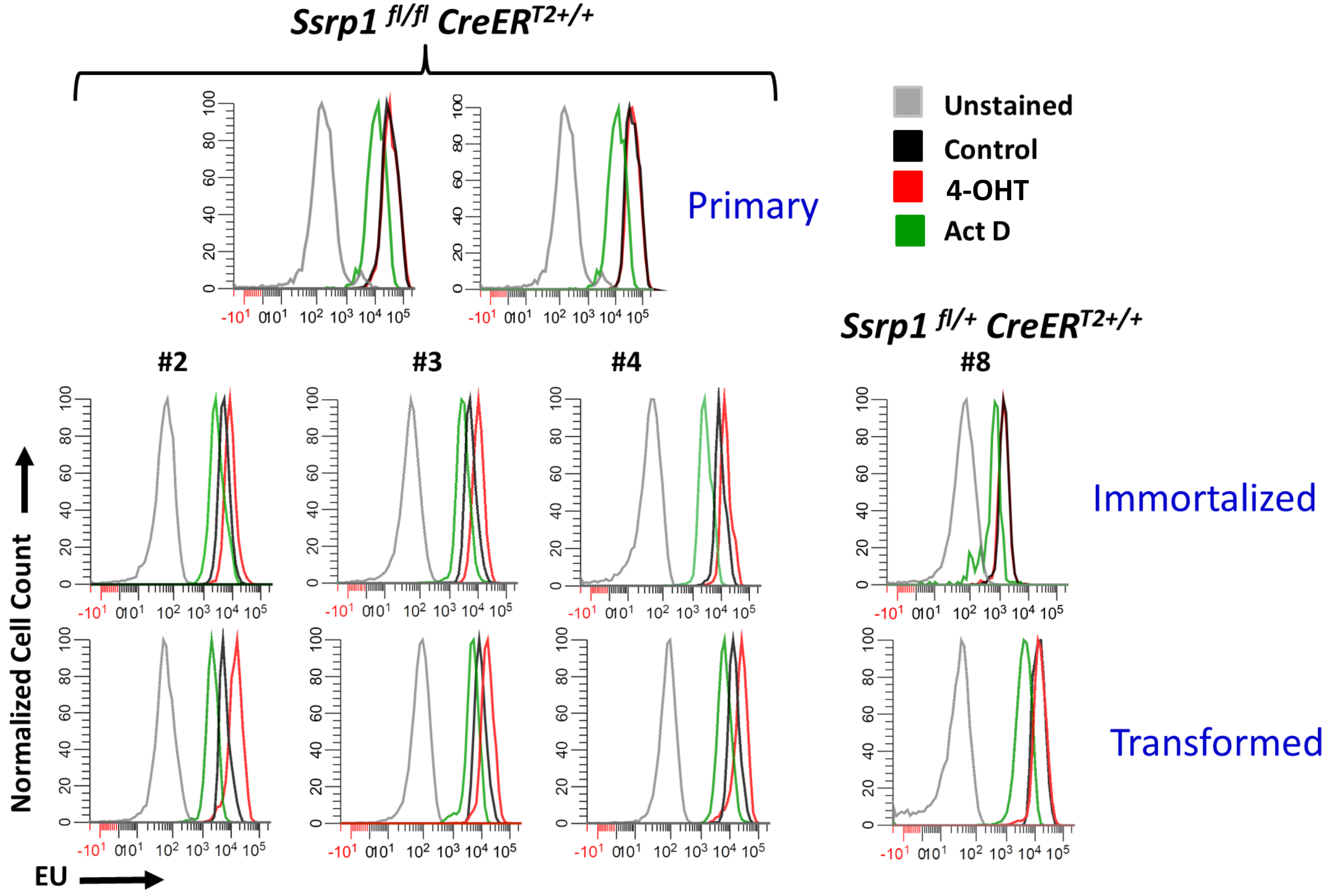


**Figure S7. Effect of *Ssrp1* KO on EU incorporation in primary (Pr), immortalized (Im), and transformed (Tr) cells.** Cells were treated for 15 min with EU 24 h after the end of the 5-day 4-OHT treatment. EU incorporation in control (black), 4-OHT (red) and Actinomycin D (ActD; green-treated Pr, Im, and Tr *Ssrp1*$^{fl/fl}$ *CreER*$^{T2+/+}$ (#2, #3, #4) and *Ssrp1*$^{fl/+}$ *CreER*$^{T2+/+}$ (#8) fibroblasts was measured using fluorescent activating cell sorting.
