## Supplementary material for "Prevention of chromatin destabilization by FACT is crucial for malignant transformation"

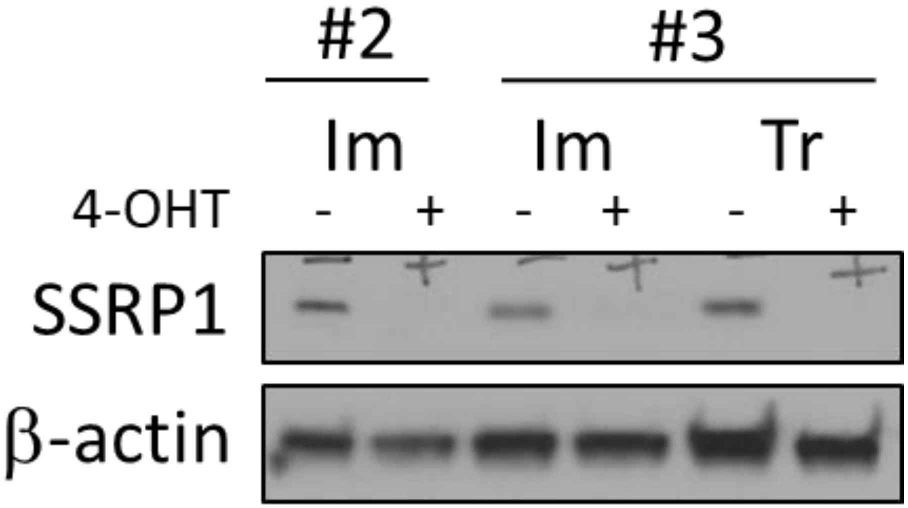


**Figure S6. Excision of Ssrp1 from growth arrested cells.** Cells were grown to confluency and then medium was replaced with serum-free medium containing 2 μM 4-OHT or vehicle. Five days after the start of treatment, cell lysates were analyzed by western blotting with indicated antibodies.
