## Supplementary material for "Prevention of chromatin destabilization by FACT is crucial for malignant transformation"

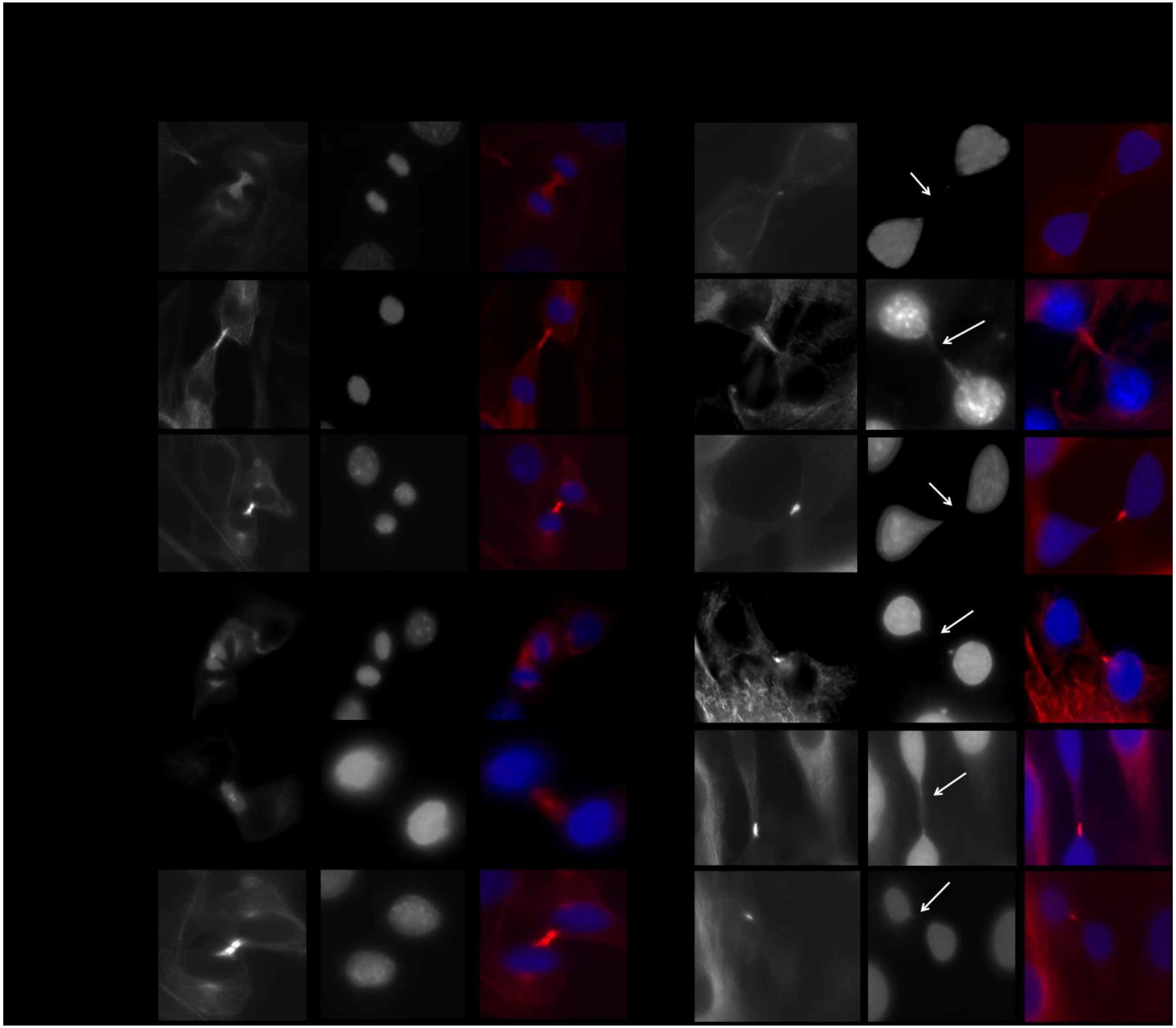

**Figure S5. Replication stress and DNA damage in immortalized (Im) and transformed (Tr) following *Ssrp1* KO.** Representative images of cells during anaphase deonstrate improper chromosome segregation following treatment with 4-OHT (white arrows) identified by staining for α- tubulin (red) and DNA (Hoechst, blue).
